# Direct single cell observation of a key *E. coli* cell cycle oscillator

**DOI:** 10.1101/2023.03.30.533363

**Authors:** Ilaria Iuliani, Gladys Mbemba, Marco Cosentino Lagomarsino, Bianca Sclavi

## Abstract

A long-standing hypothesis sees DNA replication control in *E. coli* as a central cell cycle oscillator at whose core is the DnaA protein. The consensus is that the activity of the DnaA protein, which is dependent on its nucleotide bound state, is an effector of initiation of DNA replication and a sensor of cell size. However, while several processes are known to regulate DnaA activity as a function of the cell cycle, the oscillations in DnaA expression and DnaA activity have never been observed at the single cell level, and their correlation with cell volume has yet to be established. In this study, we measured the volume-specific production rate of a reporter protein under control of the *dnaA*P2 promoter in single cells. By a careful dissection of the effects of DnaA-ATP-and SeqA-dependent regulation, two distinct cell cycle oscillators emerge. The first oscillator, driven by gene dosage, DnaA activity and SeqA repression oscillates synchronously, and shows a causal relationship, with cell size and divisions, similarly to initiation events. The second one, a reporter of dosage and DnaA activity only, is strongly coupled to cell size, but loses the synchrony and causality properties, suggesting that DnaA activity peaks do not correspond directly to initiation events. These findings suggest that while transcription regulation by DnaA activity performs volume sensing, transient inhibition of gene expression by SeqA following replication fork passage keeps DnaA activity oscillations in phase with initiation events.

---

The DnaA protein is a key factor for the initiation of DNA replication and an essential protein for most known bacteria ^1^. When it is in its ATP-bound (“active”) form, DnaA binds to a set of specific sites at the origin of DNA replication (oriC) leading to the formation of an oligomeric structure and the subsequent melting of an AT-rich region required for the assembly of the DNA replication forks ^2–5^. Based on population measurements, it is believed that the amount of DnaA-ATP needs to reach a threshold value once per cell cycle for this structure to form, leading to the initiation of DNA replication ^6–8^. Experiments in bulk exploring a large range of growth rates have shown that cell size at initiation of DNA replication is related to the growth rate and is correlated with the concentration of the DnaA protein ^9^. In *E. coli*, the DnaA-dependent regulatory circuit is made of different positive and negative components ^10^. Several factors contribute to the decrease in DnaA activity after initiation has taken place ^11^. Firstly, the expression of the *dnaA* gene is prevented for a fraction of the cell cycle by the SeqA protein binding to hemi-methylated GATC sites at the promoter, and within the *dnaA* gene itself, after their replication ^12–16^. The *dnaA* gene is located close to the replication origin, and SeqA follows the forks, transiently repressing its expression immediately after each initiation. Secondly, inhibition of transcription initiation by the oligomerisation of DnaA-ATP itself decreases the production of DnaA ^17–20^. This is thought to occur at the time of initiation, when DnaA-ATP concentration is at its peak. Both of these processes inhibit DnaA protein expression and are related to the timing of initiation of DNA replication and to fork progression through the genomic position of its gene ^13^. After initiation has taken place, the rate of hydrolysis of the ATP bound to DnaA is increased via an interaction with the Hda protein mediated by the *β*-clamp during ongoing DNA replication in a process called RIDA (Regulatory Inactivation of DnaA) ^21^. Finally, the binding of DnaA-ATP to the *datA* site also contributes to the conversion of DnaA-ATP to DnaA-ADP after initiation, via *datA*-dependent DnaA-ATP hydrolysis (DDAH) ^22^. The increase in DnaA-ATP required for the initiation of a new DNA replication cycle depends on the timely accumulation of newly expressed protein ^7, 8, 13^ and on the binding of DnaA to the DARS1 and DARS2 sites, contributing to a further increase in the DnaA-ATP to DnaA-ADP ratio by favoring the exchange of the DnaA-bound nucleotide ^10, 23–25^.

Together, these processes lead to the belief that DnaA activity is a cell cycle oscillator. Specifically, oscillations in DnaA activity are believed to play a key role at faster growth rates, when the frequency of initiation increases relative to the time of genome replication ^12, 26–28^. How-ever, given the complexity of this regulatory circuit, one of the major challenges in the field has been to find a way to measure the oscillations in DnaA activity in real time, particularly because they occur at the level of the single cell and they are not synchronized across cells in a normally dividing population. Moreover, while the different mechanisms regulating DnaA activity via ATP hydrolysis have been carefully characterized, less is known on the real time dynamics of DnaA’s gene expression as a function of the cell cycle and its coupling to cell size. In this study, we rely on a chromosomal promoter-reporter system based on the *dnaA* promoter itself as a reporter of the relative contributions of DnaA-ATP and SeqA on the expression of DnaA and to measure their relationship with cell size and cell division.

Monitoring these oscillations in single cells is particularly important because it makes it possible to compare the oscillator with known single cell observations of replication-initiation reporters ^29–36^. These studies reported a constant (independent from cell size at initiation) added size between initiation events and also between the initiation of DNA replication has and cell division ^33^. However, a direct link is still missing between possible single cell oscillations in DnaA activity and cell size correlation patterns of initiation events.

## The *dnaA* promoter as a reporter of the changes in DnaA-ATP activity and SeqA *in vivo*

Since DnaA in the cell exists under two forms, ATP and ADP bound, only the former being the one that can initiate DNA replication, we looked for a reporter of DnaA-ATP activity to study the regulation of DNA replication in *E. coli* in real time *in vivo*. We have chosen to use the role of DnaA as a transcription factor ^39, 40^ to report on the changes in DnaA activity. One of the best characterized targets of transcription regulation by DnaA is its own promoter ^18, 19^. We have constructed a reporter cassette where the fast-folding *mut2-gfp* gene is under control of the *dnaA*P2 promoter sequence (from -136 to +48 relative to the transcription start site). This GFP protein is highly soluble and stable ^41^. This construct includes a Kanamycin resistance cassette expressed divergently upstream from the *dnaA* promoter. In order to obtain an effect of SeqA and gene dosage on our reporter as similar as possible to the endogenous promoter, we have inserted the *dnaA*P2 promoter reporter cassette in the genome within the Ori macrodomain, at the ”Ori3” locus downstream of the *aidB* gene (4413507 bp) ^37^, which was used in a previous study ^42^. The coordinate of this site is at 21% of the right replicore, the replication fork should pass through it on average about 8 minutes after initiation of DNA replication.

Regulation of expression of the *dnaA* gene depends on a promoter region that includes two promoters, P1 and P2 ^43^. P2 is found downstream of P1 and includes a GC-rich discriminator region overlapping with the transcription start site, making transcription initiation negatively regulated by ppGpp ^44^. P2 is the stronger promoter in exponential phase and is thought to provide the main growth-rate-dependent regulation of DnaA expression, while P1 provides a basal level of constitutive expression, similarly to what is found at ribosomal promoter regions ^19, 44, 45^.

Expression from P2 is negatively regulated by a high concentration of DnaA-ATP, and positively regulated by DnaA when its concentration decreases ^18, 19, 46^, making it an effective sensor of DnaA-ATP levels. More specifically, two high-affinity sites for DnaA, Box1 and Box2, are found upstream of the *dnaA*P2 promoter. The binding of DnaA-ATP to these two high-affinity sites activates transcription when DnaA-ATP activity is low. As DnaA-ATP concentration increases, the DnaA bound to Box1 and 2 becomes the scaffold for the formation of an oligomeric structure that represses transcription by occluding the RNA polymerase binding site ^18, 19, 46^. Specific mutations in Box1 and Box2 disrupt the DnaA-binding consensus sequence. These mutants decrease the binding affinity for DnaA and thus result in promoters that are only positively regulated (Box1 mutation) or neither positively nor negatively regulated by DnaA-ATP (Box1-Box2 mutation) ^19^. Finally, transcription initiation is inhibited by SeqA binding to five GATC sites, two overlapping with the -10 and -35 sequences of P2 and three closely spaced sites found downstream of the transcription start site.

An additional set of mutations (“m3SeqA”, Fig. 1A) change the sequence of these three GATC sites. A previous study has shown that the set of these three mutations does not affect the synchrony of initiation of DNA replication, but causes a decrease in growth rate and DNA content in rich media^27^. The two GATC sites overlapping with the -10 and -35 sequence were left intact in order not to affect the binding of RNA polymerase, but a previous study has shown that they have little effect on repression by SeqA and DNA replication parameters^47^. As a further reference, we considered the expression of the same reporter fluorescent protein under control of a constitutive promoter used in a previous study ^42^. This phage-derived constitutive promoter, “P5” in Fig. 1A, has consensus -10 and -35 sequences and lacks regulation by specific transcription factors, therefore its GFP production rate can be considered to be largely representative of the change in gene copy number with the passage of the DNA replication fork. To verify that its expression depends on the change in gene copy number as a function of the cell cycle we have inserted the P5-*gfp* construct at the same origin-proximal locus as the *dnaA*P2-GFP construct (Ori3) as well as at the terminus-proximal locus (Ter3) downstream of the *uspE* gene (see Methods).

**Figure 1:**
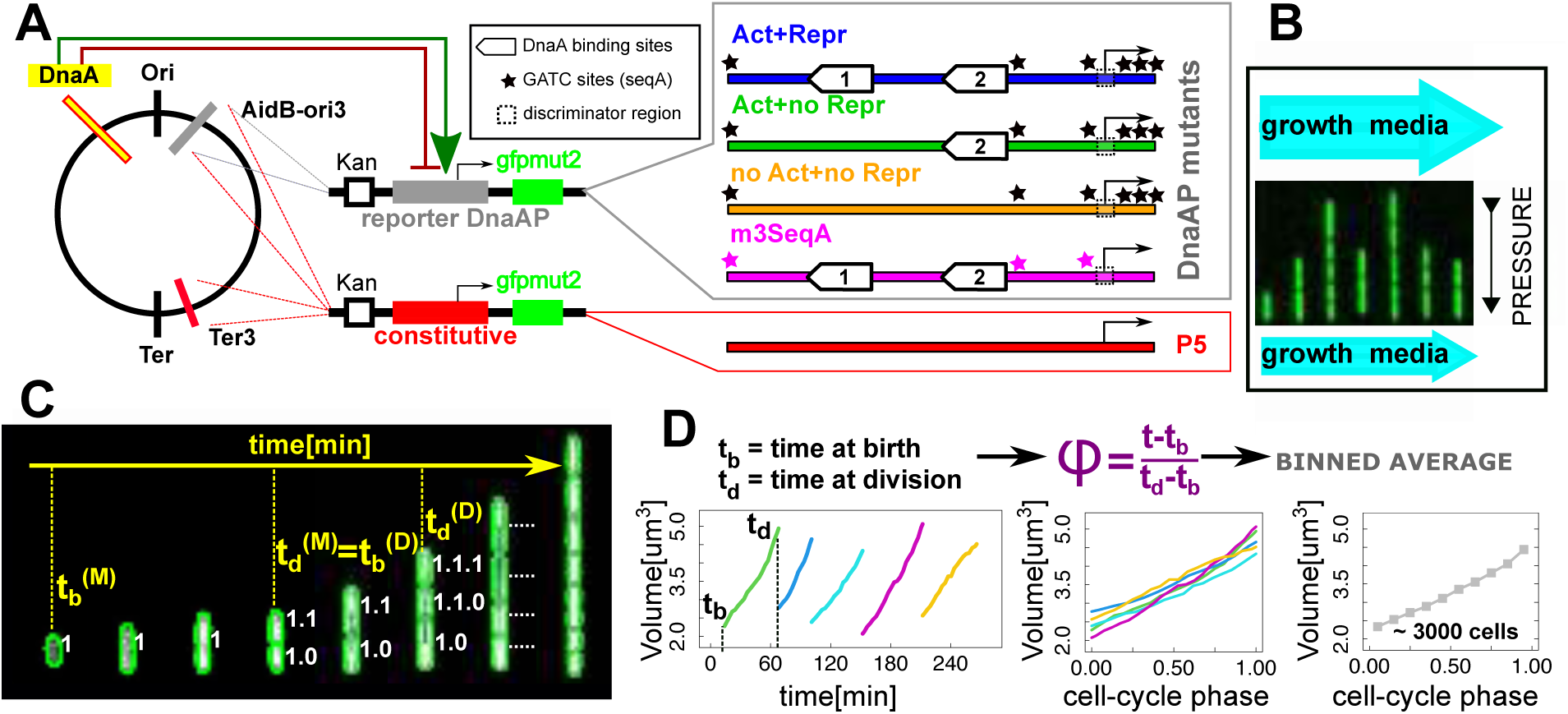
Robust and long-term single-cell tracking in fast growth conditions. **(A)** We inserted a reporter construct of the *mut2gfp* gene at a specific site downstream of the *aidB* gene in the Ori macrodomain ^37^. Expression of the reporter protein is under control of the native *dnaA*P2 promoter region. A Kanamycin resistance cassette is expressed divergently upstream of the promoter. Different promoter mutants with different levels of regulation by DnaA and SeqA were considered. As a reference for baseline gene expression we used a constitutive promoter (see Methods). **(B)** The experimental device is a two-ended ”mother machine” microfluidic channel where growth media flows constantly at the top and bottom of tapered-end micro-channels, ensuring a constant environment ^38^. The picture shows a field of view with 7 micro-channels. Differences in flow rates between the two large channels generate a pressure that keeps the bacteria (which can only escape from the larger top end) inside the channels. **(C)** Segmentation/tracking algorithms follow the changes in cell size and fluorescence over time and across generations. The image shows snapshots of the same channel at subsequent times, and *t_b,d_* represent the times of birth/division of mother/daughter (M,D) in a lineage. **(D)** To examine the effects of cell cycle progression, we aligned growth and gene expression data with respect to cell cycle phase (fraction of the cell cycle), defined as cell cycle time normalized by the cell’s division time.

The effect of the *dnaA*P2 promoter mutations on GFP expression from the chromosomal insertion site have been measured by a plate reader assay and are consistent with the previously published results obtained on a plasmid ^19^ (Supplementary Fig. S1). Mutation of Box1 increases expression relative to the wild type sequence, while mutation of both Box1 and 2 decreases expression back to wild type levels. What is important for this study is that (i) the original promoter (“Act+Repr” in Fig. 1A) senses both DnaA-ATP levels and the transit of the replication forks via the negative effect of SeqA binding (ii) the promoter stripped of both DnaA binding sites (“noAct+noRepr” in Fig. 1A) only senses the binding of SeqA, and (iii) the m3SeqA variant without the three GATC sites is only regulated by DnaA-ATP levels.

To achieve single cell resolution and capture the dynamic changes in cell growth, cell division and gene expression as a function of the cell cycle as well as across several generations, we used an integrated microfluidics and time-lapse microscopy approach ^38^. In this device, an air pressure-controlled flow system provides a constant environment where cells can grow steadily for several days as the growth medium flows continuously at a constant rate (Fig. 1B and Supplementary Fig. S2). We studied cells in a fast growth condition (M9 minimal medium with glucose and casamino acids at 30°C), where the cells have a mean doubling time of 45 minutes and initiate DNA replication at 2 origins. Thousands of single cells were segmented and tracked from movies with frames obtained every 3 minutes to examine cell cycle dependent changes in fluorescent protein expression and cell size in lineages comprising up to 15 generations, as described in ^42^ (Fig. 1C). Supplementary Table S1 provides a complete list of measured parameters and computed variables. Each experiment yielded 2-8000 full cell cycles with a good reproducibility (Supplementary Fig. S3).

## In absence of transcription regulation, GFP production rate increases with gene copy number and cell volume

To establish a solid reference for monitoring the cell cycle dependence of gene expression from the *dnaA*P2 promoter, our first goal was to characterize the “null” relationships between cell cycle progression and gene expression, i.e., the cell cycle variability of an unregulated promoter. To estimate the promoter activity in single cells, we defined a GFP production rate as the discrete time-derivative of fluorescence *dF/dt* from the time series of total fluorescence *F* (*t*). Protein production rate can vary along the cell cycle because the replication of a gene at a specific moment in time doubles the probability that it will be expressed. Since gene replication occurs at a time in the cell cycle that depends on the gene’s distance from the origin of DNA replication, the cell cycle dependence of its expression rate will depend on the gene’s location along the genome. The copy number of a gene, *g*, therefore doubles during the cell cycle, and its timing can be estimated quantitatively by a standard model ^48^, which also takes into account the case of overlapping DNA replication rounds, where replication forks from different initiation events are active in the same cell (see Methods). Additionally, a protein’s production rate has been observed to be proportional to cell size ^49^, probably in connection with the fact that cell size tends to be proportional to ribosome amounts. In our data, cell volume was computed considering a cell as a cylinder with two hemispherical caps where the radius was estimated from the segmented projected area (see Methods).

Supplementary Fig. S4A reports scatter plots of GFP production rate versus volume for origin-and terminus-proximal P5 constructs. These single-cell data show an average linear proportionality between the GFP production rate and cell volume, leading to an increase along the cell cycle. As expected from the estimation of average gene copy number, the same unregulated promoter shows an increased production rate when it is placed close to the replication origin compared to when it is found near the terminus. Normalizing the data by the estimated mean gene copy number (Eq. 3) removes most of this offset (Supplementary Fig. S4B), but the volume dependency remains.

In order to quantify the change in gene expression due to the increase in gene copy number as a function of cell cycle progression, we averaged the same data conditioning by cell cycle phase ^50^, defined as cell cycle time rescaled by the time between two consecutive divisions, i.e., *t/τ* (Supplementary Fig. S4AB). This procedure makes it possible to average together cells with all doubling times (3000-8000 cells in our case), hence increasing the statistical power. We found that the GFP production rate from the constitutive promoter is biphasic, which appears more clearly when the promoter is inserted close to the replication origin (Supplementary Fig. S4B). The beginning of the second phase correlates with the expected value of the cell cycle phase at which the gene is copied by DNA replication, supporting an effect of gene dosage. Rescaling the GFP production rate by cell volume and mean gene copy number highlights measurable oscillations that are presumably due to cell cycle dependent gene dosage (Supplementary Fig. S4C). In particular, both the ori-proximal and the ter-proximal reporters increase their production after they are replicated. These small changes can be observed thanks to the large number of cell cycles analysed in our data.

In summary, GFP production from an unregulated promoter involves “null” cell cycle dependent trends due to volume and gene dosage, which in a regulated promoter have to be disentangled from any regulatory signal.

## Transcription regulation by DnaA-ATP and SeqA causes strong oscillations in volume-specific GFP production rate

We next set out to ask how the expression of GFP under control of the *dnaA*P2 promoter differs from that of a constitutively expressed gene as a function of the cell cycle. There is only a small reproducible difference in the concentration of GFP as a function of the cell cycle phase when the constitutive and *dnaA*P2 promoters are compared (Fig. 2A). However, the volume-specific GFP production rate from *dnaA*P2 clearly shows an oscillation that is not present in the data for the constitutive P5 promoter or the promoter mutant without DnaA-ATP regulation (“no Act+no Repr”, with mutated DnaA Box1 and 2 binding sites in Fig. 1A), which behave similarly in these conditional averages, despite of the repression from SeqA and the different promoter se-quences (Fig. 2B). Using lineages of two generations we also tested whether these cell cycle phase oscillations are detectable in mother-daughter lineages, which is indeed the case (Fig. 2C).

**Figure 2:**
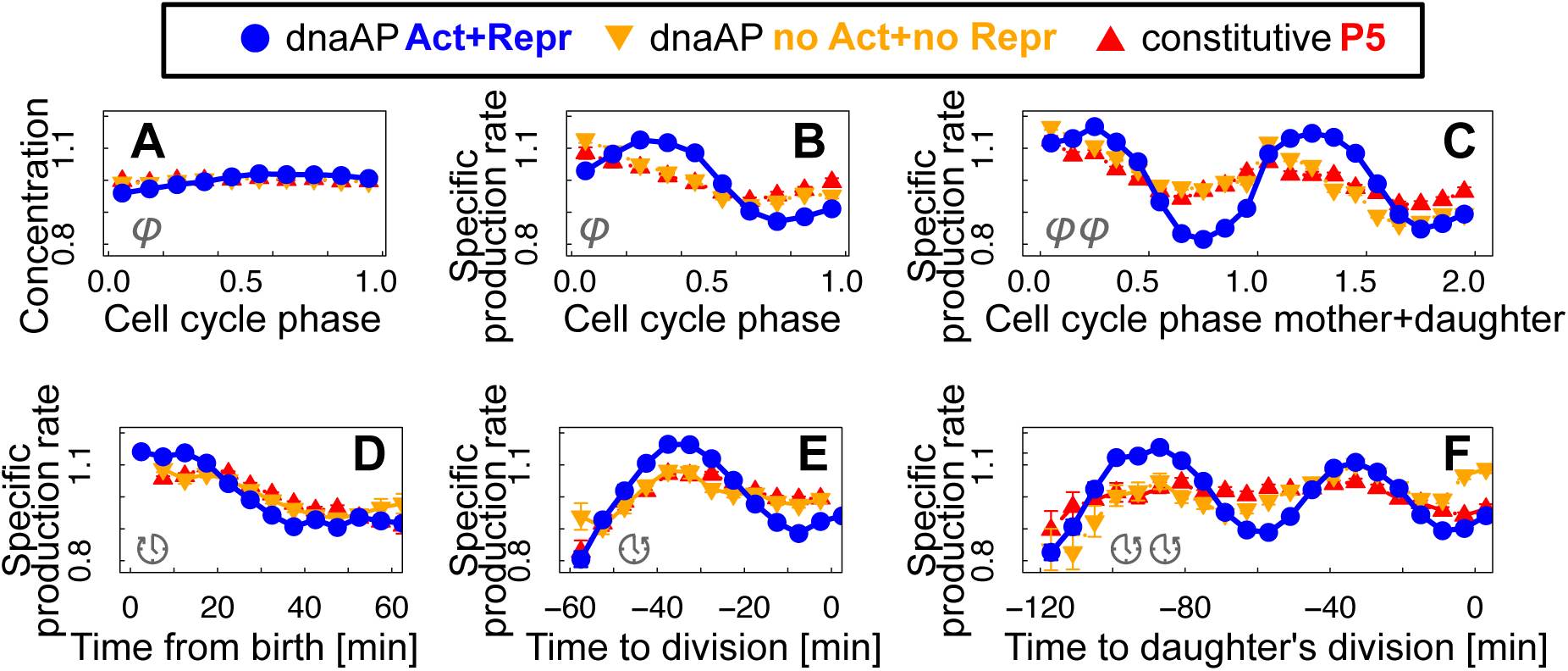
Regulation of the *dnaA*P2 promoter by DnaA-ATP causes an oscillation in GFP production rate beyond the effect of gene dosage. **(A)** Oscillations in GFP concentration as a function of cell cycle phase are weak for the constitutive promoter (red triangles), the *dnaA*P2 promoter (blue circles) and *dnaA*P2- Box1-Box2, the promoter not regulated by DnaA (orange triangles). **(B)** The fold-change in volume-specific GFP production rate from *dnaA*P2 shows a clear peak that is not present for a constitutive promoter or the *dnaA*P2-Box1-Box2 promoter, which follow similar weak trends. **C** A sinusoidal oscillation for GFP expression from *dnaA*P2 is observed when taking into account the cell cycle phase of lineages with two consecutive generations. **(D,E)** Volume-specific GFP production rate averaged conditionally to time from birth is very similar for the three promoters, while oscillations are enhanced for the *dnaA*P2 promoter when the same data are averaged conditionally as a function of the time to division. **(F)** Oscillations in GFP expression from *dnaA*P2 are enhanced when averaged conditionally to time to daughter division. If these oscillations correspond to changes in DnaA activity, they are consistent with the model where DNA replication initiating in the mother cell and terminating in the daughter cell will influence the timing of cell division of the daughter ^48^. Error bars (often smaller than symbol size) are standard errors of the mean from a re-sampled distribution obtained by bootstrapping from the experimental data for each bin.

The near-equivalence of the promoter stripped of DnaA regulation and the constitutive one in Fig. 2BC suggests that the cell cycle dependent SeqA repression alone does not suffice in establishing the reported oscillations, while DnaA-ATP oscillations are essential. The *dnaA*P2 promoter includes a GC-rich discriminator region at the transcription start site which makes it a target for both ppGpp and negative supercoiling dependent regulation ^45^. ppGpp levels and negative super-coiling have a strong effect on the regulation of DNA replication as a function of growth rate ^51^. As a control, we measured the expression of GFP from a minimal ribosomal promoter (*rrnB*P1) that also contains a GC-rich discriminator region. Expression of GFP from this promoter does not show a cell cycle dependent oscillation beyond dosage effects (Supplementary Fig. S5), confirming that ppGpp levels and DNA supercoiling are not sufficient to create the observed volume-dependent oscillation.

We can gain more insight on the relationship between these oscillators and the cell cycle by performing averages of the volume-specific production rate that are conditioned on specific cell cycle variables. Specifically, by averaging the data as a function of the time from cell birth the specific GFP production rate from *dnaA*P2 becomes almost indistinguishable from that of the constitutive promoter (Fig. 2D). However, the amplitude of the oscillation is much greater than that of the constitutive promoter once we average the data as a function of the time to division (Fig. 2E). The two findings suggest that the wild type *dnaA*P2 oscillations are somewhat agnostic of cell birth (or not synchronized with it), and prognostic of (or synchronized with) the next cell division. Fig. 2F shows that the difference between *dnaA*P2 and the constitutive control promoter P5 becomes more evident when the time to daughter’s division is used to bin the data. The reason for this is simple if we assume that minima of this oscillator can correspond to the effect of the increase in gene copy number, and thus initiations of DNA replication, amplified by DnaA-dependent regulation. The replication cycle spans two cell cycles under these growth conditions, therefore the time of initiation within the mother cell can have an effect on the time of division of the daughter cell. The results over two-generations lineages also show more clearly how oscillations in volume-specific GFP production rate are symmetric and sinusoidal when under control of the *dnaA*P2 promoter.

Once again, the oscillations are also lost for the construct where both the DnaA binding sites have been mutated (“no Act + no Repr”, orange triangles in Fig. 2DEF). However, in the m3SeqA promoter, when the three downstream SeqA sites are mutated in the context of the wild type *dnaA*P2 sequence, the oscillations do not disappear, but they change in timing and amplitude (Supplementary Fig. S6). This result is in accordance with the idea that SeqA repression of transcription creates a delay in GFP production after the gene has been copied by DNA replication and before its expression is increased by the doubling of gene copy number. The residual oscillation measured by GFP production rate from the m3SeqA promoter should be prominently driven by changes in DnaA activity, on top of dosage changes due to replication and cell division.

These results show that volume-specific GFP production rate from the *dnaA*P2 promoter is a *bona fide* cell cycle oscillator, coupled to the cell division event, and driven by DnaA-ATP levels and SeqA repression. SeqA-dependent repression plays a role in the oscillations but, alone, is insufficient to drive it.

## Oscillations in *dnaA*P2 promoter activity are a cell size sensor

The results in Figure 2 show that oscillations emerge when lineages are aligned by cell cycle phase or time to divisions, but not when they are aligned with respect to time from birth. Cell cycle phase aligns the data from individual cells as a function of the the fractional cell cycle progression between the times of cell birth and division and it has been shown to align the time of gene doubling by DNA replication ^50^. Given the consensus on the links between cell size and replication initiation, the *dnaA*P2 oscillator can be also expected to follow cell size. If that were the case, binning the gene expression data as a function of cell volume should result in an improved synchronization of individual cells’ oscillations.

We thus proceeded to test the hypothesis that DnaA activity and/or production rate may be a cell size sensor by binning the data as a function of cell volume instead of cell cycle phase. Fig. 3A shows that volume-binned averages in GFP volume-specific production rate from *dnaA*P2 follow strong oscillations reaching maxima and minima at multiples of a characteristic volume. The same analysis with the data from the constitutive promoter and the mutant *dnaA*P2 promoters also shows oscillations, however of smaller amplitude (Supplementary Fig. S7).

**Figure 3:**
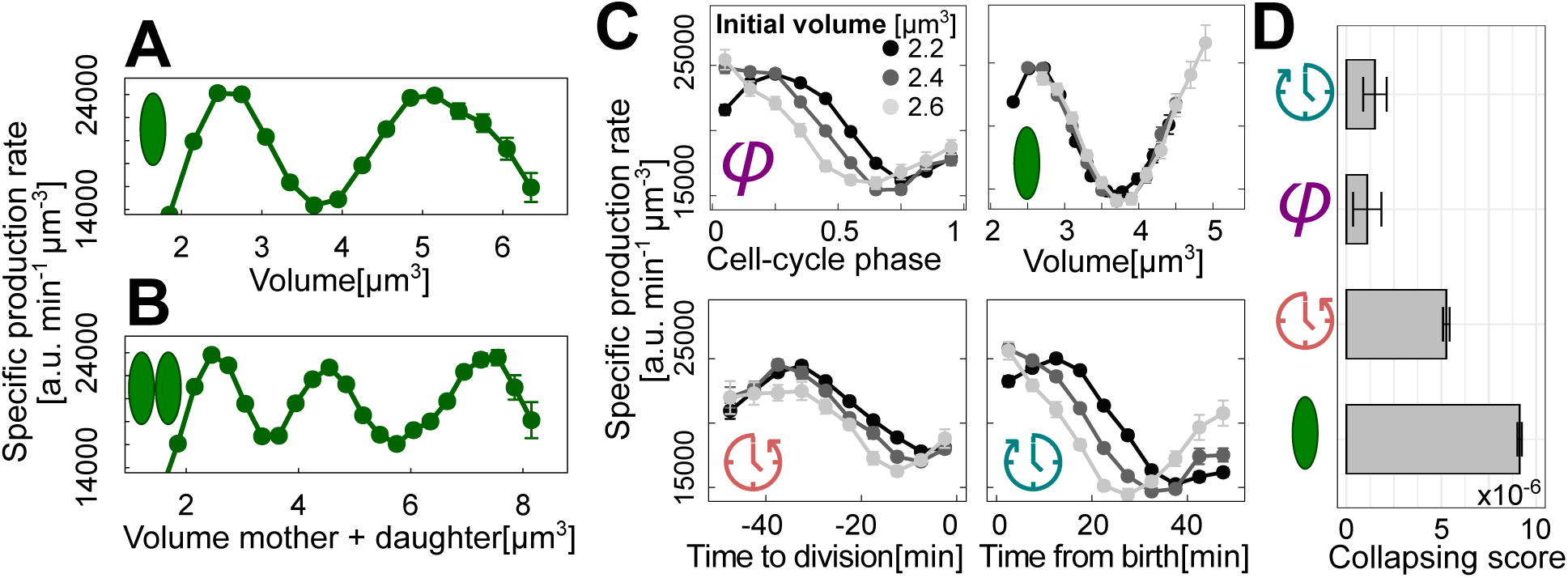
*dnaA*P2 oscillations sense cell volume. **(A)** Plot of the conditional average of GFP volume-specific production rate from the *dnaA*P2 promoter as a function of cell volume. **(B)** The same average across mother+daughter lineages shows two minima at multiples of a characteristic volume, as expected from replication initiations. **(C)** Volume-specific *dnaA*P2 promoter activity oscillations for cells of different initial size show different degrees of overlap when conditionally averaged as a function of cell cycle phase, time to division, time from birth and cell volume. The differently shaded curves result from data binned according to cell size (volume) at birth (2.2 ± 1*µ*m^2^, black, 2.4 ± 1*µ*m^2^, dark-grey and 2.6 ± 1*µ*m^2^, light-grey). If the variable in the *x* axis is the sensed variable, the averages should change independently of the cell size at birth. Data for cell volume shows the best collapse, indicating that the *dnaA*P2 oscillator is a volume sensor. **(D)** Quantification of the collapse of the curves of panel C relative to 11 bins of cell sizes at birth (see Methods). Binning the data by cell size shows the best collapse and binning the data as a function of the time to division shows a good collapse. Error bars are standard errors of the mean obtained by bootstrapping from the experimental data for each bin.

The underlying oscillation of the constitutive and unregulated *dnaA*P2 promoters reflects increased expression upon the doubling of gene dosage by the passage of the replication fork followed by a decrease as cell size further increases ^50^. Hence, part of these oscillations must be due to dosage effects and dosage-volume correlations, independently from specific regulation by DnaA or SeqA. The dosage-volume correlation is indicative of the strong cell-size dependence of initiation of DNA replication.

The effect of the mutation of the DnaA binding sites determines a visible change in amplitude and phase of the average oscillation (when averaged conditionally with respect to cell volume), consistent with a volume dependent regulation by the concentration of free DnaA-ATP, increasing gene expression rate after the increase in gene copy number and decreasing it before initiation (Supplementary Fig. S7). Thus, our data give an unprecedented view of the regulatory effect of the oscillations of DnaA activity in single cells. In the absence of DnaA-dependent regulation (noAct + noRepr) the promoter is only repressed by SeqA. In this case volume-specific production rate shows a visible change in phase compared to the constitutive promoter (Supplementary Fig. S7). On the other hand, mutation of the SeqA sites in the presence of DnaA-dependent regulation shows a visible shift in the volume at which oscillation minima occur compared to the wt *dnaA*P2 promoter (Supplementary Fig. S6). These results are consistent with repression by SeqA delaying gene expression from the newly replicated promoter.

In order to shed more light into the size-sensing properties of the *dnaA*P2 promoter, we performed joint conditional averages of volume-specific GFP production rate considering different cell cycle variables (Fig.3B). We performed these averages by further grouping cells based on their size at birth, as we figured that the variables that are more directly coupled to the oscillations should be insensitive to variations of any extrinsic variable. In particular, if the oscillator is a true volume sensor, it should have no memory of size at birth. We divided cells into 11 different birth size classes, and considered binned averages of specific GFP production rate oscillations as a function of cell cycle phase, time to division, time from birth or cell volume. Fig. 3B shows three of the birth size bins relative to an average birth size of 2.2 ± 0.1*µ*m^3^, 2.4 ± 0.1*µ*m^3^ and 2.6 ± 0.1*µ*m^3^. The plots show that volume-binned oscillations are strongly insensitive to birth size, since the data for different birth size bins overlap, while the other variables are sensitive (i.e., the plots relative to different birth size classes do not collapse). Binning by time from birth shows that cells born larger reach the minimum of the oscillation in a shorter time than cells born smaller. Binning by time to division shows the same trend, but results in an intermediate level of collapse. To quantify this behavior, we defined a score of the collapse of different birth-size classes as the inverse of the sum of SE-normalized distances between the oscillations in specific production rate for all 11 birth-size bins (Fig. 3C). The higher the score, the greater the collapse of the oscillations for cells with a different birth size. Fig. 3C shows that cell volume gives the highest score. Time to division gives an intermediate score (still a factor of two higher than those for cell cycle phase and time from birth). Finally, we compared the collapse score using different proxies of cell size (length, surface, volume) as candidates to be sensed by the *dnaA*P2 oscillator, finding that volume is the best candidate ^52^ (Supplementary Fig. S8).

## Volume-sensitive *dnaA*P2 oscillations require activation and repression by DnaA-ATP and are linked to cell division via repression by SeqA

As noted above, the average specific GFP production rate binned as a function of cell volume for the constitutive promoter, the promoters stripped of DnaA-ATP binding sites and the m3SeqA promoter show oscillations that are lower than the wild-type *dnaA*P2 promoter, but still have an amplitude of 30-40% (Supplementary Fig. S7). Crucially, however, the analysis on the joint conditional averages as a function of birth volume reveals direct volume sensing more effectively (Supplementary Fig. S9).

The comparison of the results of this volume-sensing analysis obtained with the different promoter constructs shows that the collapse score for volume is the highest for the wild-type *dnaA*P2 sequence, remains high for the mutants that are still activated by DnaA and is lost when both DnaA binding sites are mutated (Supplementary Fig. S9). Removing cell-cycle dependent repression by mutating the SeqA regulatory sites (m3SeqA reporter mutant) has a small effect on the collapse with volume but loses the collapse with the time to division (Supplementary Fig. S10). In other words, regulation by DnaA-ATP alone is not sufficient to link the oscillation in GFP production rate to the time of cell division, which seems to require repression by SeqA.

Finally, regulation by SeqA alone (in the mutant promoter not regulated by DnaA) is insufficient to couple the oscillations to either volume or cell size at division independently of birth size (Supplementary Fig. S9). Note that the population average gene expression level for this mutant is very similar to that of the wild type (Supplementary Fig. S1). In summary, both positive and negative regulation by DnaA are required to maintain a good coordination between the oscillator and cell volume. An improved level of collapse of the curves is obtained by transcription repression from SeqA.

Together, these data lead us to conclude that the *dnaA*P2 oscillator sets its phase through the timing of SeqA regulation and the changes in gene copy number. These oscillations are amplified by volume-coupled DnaA-dependent transcription regulation due to the cycle of DnaA activity.

## At the single cell level, *dnaA*P2 oscillation minima are compatible with initiations

While all the above analyses firmly establish the coupling of the *dnaA*P2 oscillator with cell size and cell division using conditional means, more can be learned considering behavior of single cells ^53^. It is important to realize that when using conditional (binned) averages, if some oscillations exist but are not synchronized with respect to the conditioning variable, they cancel out. In other words, a flat conditional average results both if the oscillations are not there and when they are present but not in phase with each other. Instead, direct analysis of single cell lineages resolves this ambiguity. In particular, the correlation signatures linking the key events of DNA replication and cell division are well known ^30, 32, 33^, and our data allows us in principle to ask how single cell *dnaA*P2 and m3SeqA oscillations relate to cell cycle progression.

Fig. 4A and Supplementary Fig. S11 show that in our data *dnaA*P2 oscillations are detectable at the level of a single cell track. We defined an automated algorithm that extracts for every cell and lineage the local minima of the *dnaA*P2 oscillations. The procedure to detect the minima is delicate, as it requires taking derivatives of noisy data and smoothing, hence subject to false positives (see Methods). Despite this limitation, our data gave us the cell cycle events related to *dnaA*P2, plus cell division, for each cell and lineage, where we know instantaneous cell size, growth rate, and interdivision time of each cell. Our data lack a direct proxy for the initiation of DNA replication, but we could ask whether the *dnaA*P2 oscillations minima in single cells follow similar patterns to the ones recently observed for initiation of DNA replication ^29, 32, 33^. We focus in particular on the minima, which could occur downstream of initiation of DNA replication, the moment where SeqA represses transcription and DnaA-ATP is near its highest activity.

**Figure 4:**
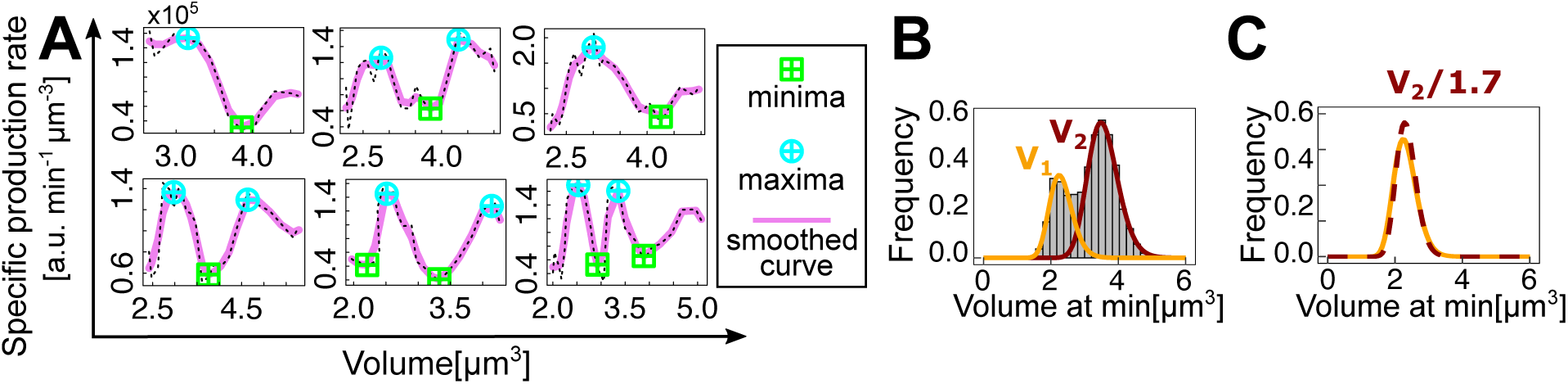
In single cells, minima and maxima of *dnaA*P2 oscillations appear at multiples of a characteristic volume similarly to DNA replication initiations. **(A)** Oscillations are detectable at the level of single cells. The plots show some single cells *dnaA*P2 tracks *vs* volume (dashed lines), as well as the smoothed tracks (purple solid lines) and the local maxima and minima (symbols) found by our automated algorithm (see Methods). **(B)** The distribution of cell sizes at minima is bimodal with peaks at discrete cell volumes. The bar plots show histograms of cell size at *dnaA*P2 minima, and the solid lines are a fit with a log-normal mixture model, which results in two log-normal distributions. **(C)** If the minima are consistent with initiations, the two distributions should collapse if we divide the mean of the second peak by 2. The best collapse is obtained for a factor of 1.7.

In the fast growth condition of our experiments, we found that the distribution of cell sizes at minima is bimodal (Fig. 4B). This is because in some cases, at fast growth, initiation of DNA replication takes place before cell division has occurred, and thus at twice the number of origins and at a cell volume that should be proportional to the number of origins. This is in agreement with the fact that the distribution of cell sizes at initiation of DNA replication in the presence of overlapping rounds of DNA replication should be approximately the sum of two log-normal distributions with means that are one the double of the other ^54^. Using a log-normal mixture model to separate the two distributions, we tested whether they would collapse by dividing the mean of the second distribution by two. In our data, the best collapse is achieved by dividing by a factor of 1.7. The small discrepancy could be due to false-positives in our detected minima and to correlations between the volumes at initiation and the probability of extra rounds of replication, previously observed computationally ^54^. Additionally, the knowledge of *dnaA*P2 oscillations minima along single cell lineages allowed us to repeat the analysis reported in Fig. 3 and test for direct sensing of added volume using doubly-conditioned averaged with added volume and volume at birth. Supplementary Fig. S12 shows that the collapse of added volume is as good as for volume itself, hence the analysis does not select which of the two variables is more tightly connected with the oscillator.

Next, we decided to investigate the correlation patterns that link consecutive *dnaA*P2 oscillation minima across one cell division. As a first control, we verified that our cells show adder correlation patterns between cell birth and division independently of cell size at birth (Supplementary Fig. S13) ^55, 56^. Supplementary Fig. S14 shows that the added volume between two minima weakly correlated with cell volume at the first minimum. Considering the main mode of the distribution, the slope of the conditional average was −0.266 ±0.004 and consistent across one replicate (−0.19 ± 0.09). Considered the uncertainty due to measurement noise, derivatives, and minima detection, these results can be considered in line with the adder-like inter-initiation correlation pattern found by labeling origins or replication fork proteins in single cells ^29, 31–33, 54^.

To summarize, the minima extracted from *dnaA*P2 oscillations in single cells follow single-cell correlation patterns that are roughly in line with replication initiations, supporting our interpretation that the *dnaA*P2 cell cycle oscillator is intimately linked to the single cell replication cycle.

Performing the same analysis on the m3SeqA oscillator minima, (Supplementary Fig. S14) we found a slightly stronger slope, suggesting that the two minima follow different rules. We note however that this difference was found only in one of the two experimental replicates for m3seqA at fast growth, probably due to uncertainties in minima detection in the lower-quality dataset. This observation is mirrored by the shift of the minima at smaller volumes and cell cycle phase observed in conditionally averaged m3SeqA oscillations (Supplementary Fig. S6).

In summary, the results of the analysis at the single-cell level support those obtained from conditional averages, they indicate that the wild type *dnaA*P2 oscillator acts in synchrony with volume and with cell division, while the m3seqA oscillator, deprived of SeqA repression at fork passage, may loose some synchronization properties with the cell division cycle.

## Causal links with cell volume and division differ between *dnaA*P2 and m3seqA oscillations

Since the detection of the minima requires smoothing the data and taking derivatives, then smoothing again to detect minima reliably, this procedure is particularly sensitive to false positives due to propagated measurement noise. Additionally, the analyses so far are insufficient to hypothesize any causal links between the oscillators and the cell division cycle.

To address these problems, we used synchronization analysis, cross-covariance, and causal inference of *dnaA*P2 time series along lineages of single cells. The two latter techniques in particular have the advantage of considering the whole time series (hence leveraging all the data) and not just relying on minima detection. For each lineage, we considered the *dnaA*P2 and m3SeqA cell cycle oscillators, the cell volume time series, and the cell division time series, as three *a priori* independent signals, to investigate their synchronization and to test whether a causality between these signals exists (Fig. 5A and Supplementary Fig. S11). Each of these parameters displays some periodicity throughout several generations.

**Figure 5:**
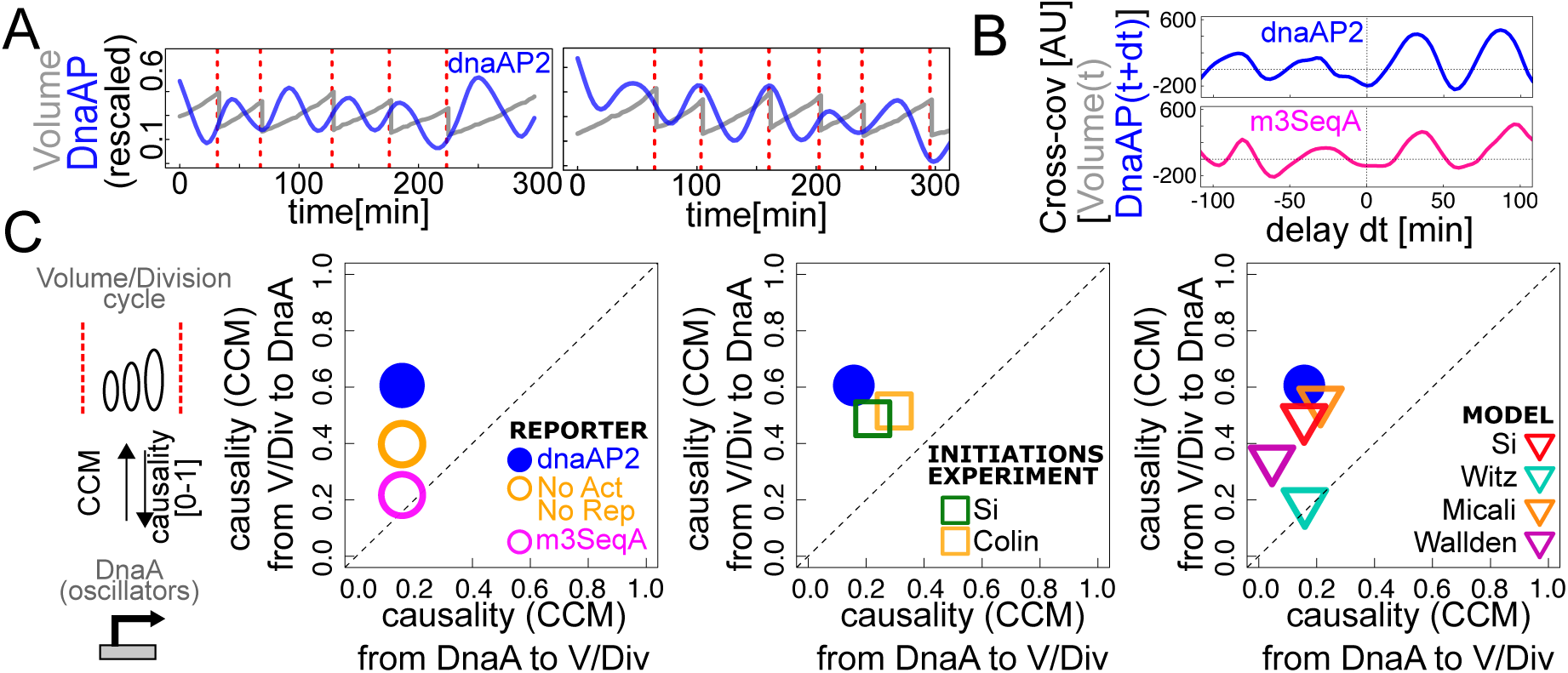
*dnaA*P2 promoter activity and growth-division are coupled oscillators. **(A)** Lineages of single cells show a clear oscillation of *dnaA*P2 promoter activity (blue), compared here to volume growth (grey) and division events (red), which we considered as three *a priori* independent time series. Volume and *dnaA*P2 promoter activity are normalized here by their average value in order to show them in the same plot. **(B)** Cross-covariance between *dnaA*P2 promoter activity and volume growth is periodic, supporting synchronization between the two oscillators, and asymmetric, supporting a stronger effect of volume changes on future *dnaA*P2 changes than *viceversa*. Asymmetry and cross-covariance are weaker for the m3SeqA oscillator. **(C)** Convergent Cross-Mapping (CCM) was used to detect causal relationships between oscillators. Left panel: For the wild-type promoter, volume/division are strong causes of *dnaA*P2 changes, but the strength of this causal link decreases in the mutants of the DnaA binding sites, and the causality becomes completely symmetric in the mutant of the SeqA binding site. Middle panel: CCM derived from experimental replication-initiation datasets is consistent with our data for the wild-type promoter. Right panel: CCM from simulated initiation datasets is most consistent with our data for the wt *dnaA*P2 promoter for the models from refs. ^29,30,32,33,35^.

To test the presence of a time hierarchy connecting specific *dnaA*P2 activity and volume os-cillations, we first considered the cross-covariance between these two time series, computed along lineages ^57^. Fig. 5B shows that this function is markedly periodic, suggesting a strong coupling, with higher-amplitude peaks for positive time delays. This asymmetry of the cross-covariance function could suggest a potential time hierarchy between volume and *dnaA*P2 whereby changes in volume may be used to predict future changes in *dnaA*P2 oscillations in volume-specific production. Importantly, in promoter mutants with mutated DnaA binding sites or in the constitutive promoter the cross-covariance are strongly reduced (Supplementary Fig. S15). Mutation of SeqA binding sites (m3seqA) mildly reduces the amplitude of the delayed cross-covariances, but alters the pattern of the time asymmetry (Fig. 5B and Supplementary Fig. S15).

Since the results on the m3SeqA promoter lead us to hypothesize that the wild-type *dnaA*P2 oscillator is in synchrony with cell cycle progression through SeqA, hence through sensing of the initiation event by fork transit, we tested the phase locking of the two oscillators as follows ^58–60^. We characterized each oscillator with a phase, a linearly increasing variable reset at each cycle, which advances by 1 between successive cycles. Specifically, we defined a *dnaA*P2 phase variable Θ, where Θ = 0 corresponds to the minima of *dnaA*P2 promoter activity and a cell cycle phase Φ as described above in this text (Φ = 0 represents birth). These two variables define a periodic square (a torus) [0, 2*π*)x[0, 2*π*). Subsequently, as time *t* increases, the combined phase (Θ,Φ) from each single cell lineage traces a trajectory in this phase space. If their orbits are phase-locked, single lineages follow a diagonal trajectory in this space. Hence, we took a heatmap of the trajectory density from all available data (Supplementary Fig. S16). For the wt *dnaA*P2 oscillator, the phase difference, Δ = Θ−Φ fluctuates around a constant value, hence the heatmap shows two juxtaposed diagonal stripes whose slope is one. Hence the oscillations only depend on the phase difference, as holds for a wide class of synchronized oscillators ^61–63^. Conversely the m3SeqA mutant breaks this dependency, likely as a result of the loss of complete synchrony with the cell cycle phase. Note however that contrary to the cross-covariance analyses, definition of a phase relies once again on delicate minima detection.

Due to the loss of synchronization with the cell cycle of the m3SeqA oscillator, and due to its loss of delay-time asymmetry in the cross-covariance with volume, we figured that the causal links between the oscillator and the other proxies of the cell cycle may be stronger for the wild-type *dnaA*P2 oscillator. While asymmetric cross-covariances reveal a time hierarchy, one has to be careful when inferring causal relations between two observables because of the existence of a non-zero correlation between two signals does not necessarily imply a causal link ^61^. To investigate the directionality of the coupling, we used the Convergent Cross Mapping (CCM) technique, considering *conditional* correlations of one variable with a second one, under the constraint that the former is constrained to its attractor manifold, as reconstructed from the time series using Takens’ theorem ^64, 65^ (see Methods and Supplementary Fig. S17). Importantly, this analysis is based on the whole time series and does not rely on minima detection. The output of this analysis, given two time series *A* and *B* is a pair of a directional parameters *ρ_AB_* and *ρ_BA_*, between 0 and 1, that represent the strength of the causality link from *A* to *B* and from *B* to *A*. A causal link is witnessed by unequal causality parameters in the two directions *ρ_AB_* ≠ *ρ_BA_*.

Fig. 5C summarizes the results of this analysis on our data. In order to try to disentangle the effects of cell volume from cell division events, we first tested the causality between cell volume and an oscillatory signal constructed to have a maximum at division. This analysis returns that volume and cell division are always in a strong symmetric relationship (Supplementary Fig. S18), hence, they are causally indistinguishable. Crucially, Fig. 5C shows instead a strongly asymmetric causality from volume (or division) to *dnaA*P2. This asymmetric causal link is weakened in the promoter without negative and positive DnaA regulation, and disappears in the m3SeqA promoter, for which the correlation becomes causally symmetric. It is important to point out that these results do not refer to the mutant of the endogenous *dnaA* promoter, but only report how the same endogenous DnaA-ATP oscillations and SeqA transit are read by our reporters when there are mutations of the binding sites. Hence, the observation that causality is lost when SeqA binding sites (and thus sensing of the passage of the replication fork) are mutated indicates once again that the endogenous *dnaA* promoter as well as our reporter takes relevant input from the fork transit.

Since we have found that *dnaA*P2 oscillatory minima in single cells follow similar patterns to those found for initiation events, we devised a way to compare the oscillator’s causality patterns with those observed for initiations, both in data from refs ^29, 32^ and in cell-cycle mathematical models proposed in the literature ^29–33, 35^ (Fig. 5C). To do this, we constructed time series connecting measured or simulated initiation events by sinusoids, in such a way that the minima coincide with initiations in time series taken from data or mathematical models (see Methods). This procedure produces a differentiable oscillatory curve, which can be compared to volume and division time series using convergent cross-mapping. Fig. 5C shows that the predicted causality pattern from experimental data on initiations is completely consistent with the one shown by *dnaA*P2. Fig. 5C shows that only the models from refs ^30, 32^ are consistent with the causal asymmetry. Both models assume that the pattern between initiations is an adder per origin. The model proposed by Si and coworkers assumes that division is agnostic of the chromosome, and not linked to replication-segregation. The concurrent processes model proposed by Micali and coworkers assumes concurrency of time scales between a cellular process and chromosome replication-segregation setting division through an AND gate.

Going back to our experimental data from wild type *dnaA*P2, our causality analysis using convergent cross mapping shows that cell division or cell size cause *dnaA*P2 oscillations in a much stronger fashion than *dnaA*P2 causes division. This perhaps unintuitive result may have two explanations, (i) there is a strong symmetric coupling between cell division and cell size, and the *dnaA*P2 promoter is a strong size sensor, or (ii) there is a strong symmetric coupling between cell division and cell size, and the *dnaA*P2 promoter is a strong cell division sensor. However, the causal equivalence of cell division and volume in our data do not allow us to distinguish between the two hypotheses. Note that given the essentiality of the SeqA binding sites for the causal asymmetry, it seems reasonable to assume that the crucial sensed event is fork transit, hence (ii) can be restated by saying that replication initiation itself may be a strong sensor of the previous cell division, and (i) that it may be a strong sensor of cell size.

In order to attempt to resolve this question, we performed additional experiments inhibiting division by adding cephalexin (Supplementary Fig. S19). Cephalexin-treated cells do not divide, but keep growing and initiating DNA replication, producing an array of nucleoids. Hence, their cell cycle is still somewhat operative. Convergent cross mapping analysis is technically impossible under this perturbation, because the volume time series increases monotonically and lacks an attractor. Interestingly this perturbation does not ablate time-periodic *dnaA*P2 oscillations (Sup-plementary Fig S19), but conditional averages become flat as soon as the cell sizes exceed the physiological range, suggesting that the synchrony with cell volume is lost ^66^.

## Discussion and Conclusions

An oscillation in DnaA-ATP activity coupled to the cell cycle in *E. coli* is assumed by most, but so far supported only by indirect population-level data ^67, 68^. Our data provide a first-time observation of a cell cycle oscillator of gene expression in single *E. coli* cells through the volume-specific production rate of a reporter protein under control of the *dnaA*P2 promoter. This promoter was chosen because it integrates, similarly to the endogenous DnaA production rate, both changes in DnaA activity, via activation and repression by DnaA-ATP, and the timing of initiation of DNA replication via SeqA repression following gene duplication.

The changes in gene dosage across the cell cycle affect the transcription rate and set up an underlying cell cycle oscillation of gene expression rate ^50, 69^. We see this in our data as a change in the volume-specific production rate from a constitutive promoter depending on its genome position (Fig. 2).

When we look at the volume-specific production rate from the *dnaA*P2 promoter and its mutants through conditional averages, we see a clear oscillation as a function of cell cycle phase as well as cell volume (Fig. 2 and 3). The oscillations have a considerably larger amplitude than those of the constitutive promoter, showing that the additional regulation of gene expression by DnaA amplifies the effect of the gene copy number. The idea that these oscillators may be strong volume (or added volume) sensors is confirmed by our analyses with doubly-conditional averages. Furthermore, changes in the volume-specific production rate measured by binning the data as a function of cell volume have a larger amplitude than conditional average taken as a function of cell-cycle phase, suggesting a stronger synchronization of the change in gene copy number with respect to cell volume than the fractional duration of the cell cycle.

The main insights in our study stem from comparing the wild type *dnaA*P2 reporter to promoter mutants of the DnaA and SeqA binding sites, allowing us to measure their effect gene expression separately. SeqA represses DnaA’s gene expression for a window of time (about 10 minutes) after the passage of the replication forks ^12^. Accordingly, the comparison of the m3SeqA mutant promoter with the wt *dnaA*P2 shows that the oscillations are shifted to smaller volumes or an earlier cell cycle phase (Supplementary Fig. S6). This also shows that direct regulation by SeqA is not required for relevant cell cycle oscillations, but regulation by changes in DnaA-ATP concentration are sufficient. The comparison of the oscillations of the constitutive promoter and the *dnaA* promoters with mutations of the DnaA binding sites is consistent with a scenario where repression by DnaA-ATP takes place mainly before the increase in gene copy number and activation by DnaA-ATP takes place after SeqA has dissociated (Supplementary Fig. S7). Connecting to standard views on this oscillator ^68, 70^ we can speculate that a rapid decrease in the free DnaA-ATP concentration after initiation is likely due to the combined effects of RIDA, DDAH, titration to newly replicated genomic sites and transient SeqA repression of DnaA expression coupled with dilution due to cell growth ^11^. Future experiments monitoring the behavior of our reporters in mutants could help elucidating these details.

Based on our analyses of conditional averages for the sensing of different variables, both the wild type *dnaA*P2 oscillator and the m3SeqA oscillator sense cell volume (Fig. 3 and Supplementary Fig. S10). Interestingly, the m3SeqA promoter, while showing a strong collapse with cell volume, shows poor collapse with time to division. Hence, while the relationship of the oscillator with volume depends on regulation by DnaA, the relationship of the oscillator with time to division appears to rely on the negative regulation by SeqA. Hence, we conclude that DnaA activity oscil-lations likely sense cell volume. Interestingly, expression from the promoter that is activated but not repressed by DnaA-ATP and repressed by SeqA retains a significant level of volume sensing and coupling with cell division time (Supplementary Fig. S9) despite an increased average expression rate (Supplementary Fig. S1). It seems that in these growth conditions timely repression by SeqA is sufficient to maintain some response to volume change in the absence of repression by DnaA-ATP. Negative regulation of gene expression by DnaA-ATP can integrate on the promoter the effects of the different regulatory factors of DnaA activity, setting an upper limit of DnaA-ATP concentration via the expression rate.

Most importantly, the high amplitude of the oscillations in the strains where GFP expression is under control of the promoters regulated by DnaA-ATP allows for an analysis of the minima at the level of single-cell lineages. Minima can be thus reliably detected in lineages of the wild type and m3SeqA *dnaA*P2 promoter constructs. In single cells, the minima of the oscillators pinpoint key cell cycle times: for the m3SeqA reporter these are related to the joint contributions of dosage oscillations and DnaA activity-based regulation, and for the wt reporter they are due to SeqA repression as well, and are expected to co-occur with fork passage a few minutes after replication initiation (the delay should be about eight minutes and the time difference between consecutive frames is three minutes in our experiments). Minima for the wt *dnaA*P2 oscillator have a resemblance with initiations ^29, 32, 33^, in that they show near-adder correlations between subsequent minima (Supplementary Fig. S14). Finally, when analyzed as a phase oscillator, only the the wt *dnaA*P2 oscillator shows strong synchronization properties with the cell cycle phase, while the m3SeqA oscillator shows a more irregular synchronization behavior (Fig. S16).

However, since these analyses relied on possibly fragile minima-detection techniques, we made use of techniques that rely on the full time series. Cross-covariance analysis confirms a difference in the relationship of wt *dnaA*P2 and m3seqA oscillations with cell cycle progression. Equally, when we detect causality by convergent cross mapping between size oscillations and the wt *dnaA*P2 oscillator, we see the same asymmetric causal pattern as for a sinusoidal oscillator based on experimental initiation events, robustly across data sets (Fig. 4), and once again this is not the case for the m3SeqA oscillator.

Hence, it appears from our data that the combination of only DnaA activity and dosage, proxied by the m3SeqA oscillator, does not encode initiations, but the *dnaA*P2 oscillator may do, and the additional SeqA repression is crucial for this by coupling the induction of gene expression to the initiation event, thus coupling DnaA-ATP production with the initiation mass of the following initiation. A further implication of these results is that peaks in DnaA-ATP concentration, defining the minima in the absence of SeqA, do not appear to correspond to initiation events. This is consistent with previous data showing that over-expression of DnaA does not have a strong effect on the initiation of DNA replication ^71, 72^ unlike under-expression, where DnaA becomes limiting^6, 8, 32, 67, 72–74^.

In the absence of direct measurements of the activity of the regulators of initiation of DNA replication, its coupling to cell size has been the object of several mathematical models ^26, 54, 68, 70, 75^. Most models predict a peak either in DnaA-ATP amount or concentration at the time of initiation. Our results provide the first evidence at the single cell level that there is an oscillation in DnaA-ATP concentration that is coupled with cell size and the cell cycle. The results obtained by the comparison of the different variants of the *dnaA*P2 promoter are consistent with a rapid decrease in DnaA activity following initiation. This takes place at the same time as repression of gene expression by SeqA, independently of the specific cell size or DnaA activity. This delay in *dnaA* gene expression further decreases DnaA-ATP levels. The tight coupling of the minima in gene expression with initiation events shows that the subsequent induction of DnaA promoter activity occurs rapidly after the dissociation of SeqA. Transient transcription repression by SeqA and titration of DnaA have been proposed to lead to steeper oscillations in DnaA activity that decrease the noise in initiation ^70^. Activation of gene expression by DnaA can further contribute to a step function in the induction of transcription prior to initiation.

Finally, while here we measured the expression of a reporter protein under control of the *dnaA* promoter, we can speculate based on our data that the expression of the actual *dnaA* gene depends on both volume sensing, via autoregulation by DnaA-ATP concentration, and sensing of a successful initiation, via SeqA repression. In a recent study, the mRNA of the *dnaA* gene was shown to oscillate with the cell size of fixed cells in a similar pattern to the one observed here for our reporter construct, with an effect of transcription shutoff by SeqA and a different phase from the one expected just from to the increase in gene dosage due to DNA replication ^69^. However, their technique does not have dynamic resolution along single-cell lineages and makes it possible to access only the relatively weak concentration oscillations shown in Fig. 2A.

In this study we have focused on using specific promoter variants that report on DnaA activity and the replication cycle in the wild type background and in the absence of perturbations. In the future, this approach can be used to more directly measure the effect of perturbations of regulators of DnaA-ATP of DnaA-ADP ratio on both the oscillations of DnaA activity and initiation of DNA replication.

## METHODS

### Strains and growth media

The experiments were carried out with the wild-type *E. coli* strain BW25113, the parent strain of the Keio collection ^76^, which has been fully sequenced ^77^. Promoter-reporter constructs were inserted in the chromosome close to the origin of replication at Ori3 (4413507 bp, in the region downstream of the converging *aidB* and *yjfN* genes). The *gfpmut2* gene, coding for a fast folding green fluorescent protein ^41^ is placed downstream of the chosen promoter sequence. A kanamycin resistance cassette (KanR) is divergently expressed upstream from the promoter region. The constitutive promoter cassette was also inserted in the chromosome close to the replication terminus, at the Ter3 site (1395706 bp, downstream of the converging *uspE* and *ynaJ* genes). Bacteria were grown overnight in M9+0.4% glucose at 30°C. Overnight cultures were diluted 500:1 in new growth medium and returned to the incubator for 3-4 h. This is important to guarantee bacteria to be in exponential phase when injected into the microfluidic device. Experiments were carried out at 30°C in M9+0.4% glucose + 0.2% casamino acids, with an average doubling time of 45 ± 5min. We verified that the different levels of expression of the GFP in the different strains do not have an effect on cell growth. Doubling time and cell size are consistent between all the mutants (Supplementary Fig. S20).

### Mother machine experiments

We used a microfluidic ”mother machine” device where the 1 micron channels are found between two large feeding channels ^78^. The bacteria are trapped in the microfluidic channel thanks to a narrower opening on one side. PDMS (Polydimethylsiloxane) devices from the mold were obtained by standard procedure and attached to a microscope slide by treatment with a plasma cleaner. Before loading bacteria into the device, each chip was treated with a solution of bovine serum albumin (BSA) to minimize bacterial interactions and binding to the glass or PDMS components. Devices were injected with around 150 *µL* of 2% BSA and incubated at 30°C for 1 h. Passivated chips were rinsed with freshly filtered medium and 1ml of bacterial culture was injected manually. Each feeding channel is coupled with a flow sensor. Using its feedback loop, we can monitor and control the flow rate in our microfluidic setup while keeping stability and responsiveness of pressure driven flows (Elveflow). This technology enables us to set up robust and long term microfluidic experiments. A home-built temperature control system is used to maintain the entire setup at 30°C. We use Nikon Inverted Microscope ECLIPSE Ti-E with 100X oil objective high, 1.4, NA (Numerical Aperture) lens, coupled with a Nikon Perfect Focus System (PFS) to rectify drift in focus. A xy motion plate is used to memorize and loop over different Regions Of Interest (ROIs) at a specified interval of time.

The camera captured 16 bit images at 512 x 512 pixel resolution with the length of one pixel equal to 0.1067*µ*m. The motorized stage and camera were programmed to cycle between at most 40 fields of view, each spanning roughly 8 microchannels, every 3 min.

### Data analysis pipeline

The data obtained are in the .nd2 format and are imported and analysed with the Fiji software. Background subtraction is performed using a 50 pixel rolling ball technique, and different positions are stored separately as a set of .tiff image files. Channels with a good number of bacteria are selected manually and stored in different folders. For segmentation and tracking, we started from codes developed by Mia Panlilio ^42^ and we added necessary modifications to optimize them for our experimental setup. Before starting, we select the experimental time window where bacteria were growing in a steady growth rate in a given growth medium. This window was defined by observing sliding averages of population interdivision time, growth rate and cell size at birth ^42^. To correct for segmentation and tracking errors we applied a set of filters. First, we considered only cells where both mother and daughter(s) were at least partially tracked. Second, we excluded division events where daughters had a volume outside of the interval 40-60% of the volume of the mother. This step aims to eliminate filamentous cells and segmentation artefacts. Third, we excluded cell cycles with interdivision times lower than 15 mins (the physiological lower limit is 15-20 min). Lastly, we considered cell cycles where initial/final volume, initial/final width and initial/final growth rate were inside the 95% tails of the distribution. Before computing the discrete derivative of fluorescence and volume (as central derivatives defined across three subsequent frames) we removed outliers (defined by subtracting a trend line by binned averages and identifying points lying more than 1.5 standard deviations from the baseline) from fluorescence and volume traces and we substituted them by linear interpolation. Minima were detected after removing outliers from the differentiated data, and smoothing with a 3-point average.

All datasets were grouped based on strain in different R objects. We performed an exploratory data analysis to check if datasets were consistent and if filters worked as desired. Custom functions were written to handle and analyse these large, non-uniform datasets. In particular, functions were written to compute the discrete derivatives and the different normalizations.

### Volume and gene dosage estimation

We calculate the volume describing a cell as a cylinder with two hemispherical caps,

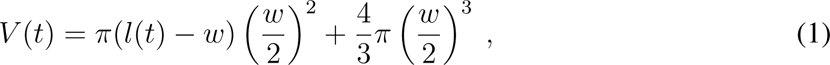

where *w* was taken as a cell cycle average of the average measured cell width. The expected gene copy number was computed from the Cooper-Helmstetter model ^79^. Replication of the *E. coli* chromosome begins from a single origin and oppositely oriented replication forks proceed symmetrically along the genome to complete replication. Since, on average, a cell divides at a time *C* + *D* (≈ 60 min) after replication initiation, an average time lag *B* before initiation is necessary to make the total replication time *B* + *C* + *D* an integer multiple of the doubling time *τ*. Thus, defining *n* = Int(*C* + *D/τ*) as the integer number of times that *τ* divides *C* + *D*, one has that *B* + *C* + *D* = (*n* + 1)*τ*. More generally, we can consider a gene at a chromosomal position defined by its normalized distance from Ori, i.e. *l* = 0 represents a gene in Ori and *l* = 1 a gene in Ter. The copy number of this gene, *g*, changes during the cell cycle following

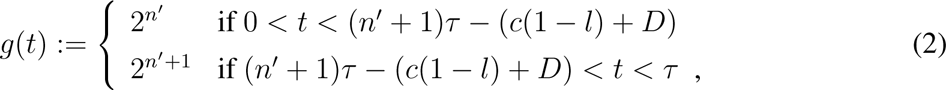

where 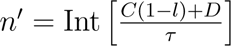. By averaging over the cell cycle one gets the expected gene

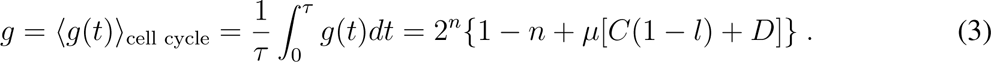

### Convergent cross mapping

Convergent Cross Mapping (CCM) is a method for causality inference based on Takens’ theorem ^64^ and developed in ref. ^65^. Takens’ theorem shows that time-delay embedding of a one-dimensional time series provides a 1–1 mapping of system dynamics from the original phase space (constructed with all system variables) to the reconstructed shadow phase space, as long as the latter has sufficient dimensions to contain the original attractor. Supplementary Figure S17 summarizes the main steps of the CCM method. Shortly, it reconstructs a shadow attractor from one time series at a time and uses these coordinates to compute conditional correlations at fixed values of the shadow attractor of the variable. Since they are based on different constraints, the conditional correlations are not symmetric, and reveal causal links. We used the R package multispatialCCM to implement CCM (https://CRAN.R-project.org/package=multispatialCCM)^80^. CCM was used on our experimental data on volume, divisions, and *dnaA*P2 oscillation time series, using DNA replication initiation data from the literature and on simulated data. In our data, we used volume and *dnaA*P2 oscillator time series, and a cell division time series was defined as a continuous process with narrow peaks around each experimental division event. For initiation of DNA replication data, we used two datasets from the literature where initiation of DNA replication and cell division were tracked within the cell cy-cle ^29, 32^, and we defined a putative *dnaA*P2 oscillator time series by assuming a sinusoid between 0 and 1 with the minima at consecutive initiations (Supplementary Fig. S17B). We also simulated four different models described in the literature: a model where replication initiation sets cell division through a timer, from ref. ^35^; a model where DNA replication initiation set cell division through an adder ^33^; a model where replication and an inter-division concurrently limit cell divi-sion ^30, 31^; and a model where DNA replication has no direct influence on the timing of division ^32^. The codes and parameter values are those used in ref. ^29^.

### Data and code availability

Data sets of segmented and tracked cells, together with example code for data analysis, were made available through the Mendeley Data Repository DOI: 10.17632/hhp6g5zt8j.1

## Acknowledgements

We are grateful to Nancy Kleckner for useful feedback on our work, and to Petra Levin, Johan Elf and Philippe Nghe for useful discussions. MCL was supported by Associazione Italiana per la Ricerca sul Cancro, AIRC IG Grant no. 23258.

