## Supplementary Fig for "Direct single cell observation of a key *E. coli* cell cycle oscillator"

### Supplementary Information for Iuliani *et al.*

### Supplementary Tables

| Notation | Name | Unit | Description |
|---|---|---|---|
| $l$ | Length | $\mu$m | Greatest pairwise distance between boundary points along the principal axis |
| $Area$ | Area | $\mu m^2$ | Area enclosed by boundary points, including edge pixels |
| $w$ | Width | $\mu$m | Width assuming the projected area is a rectangle with semicircular caps of radius $\frac{w}{2}$ |
| $V$ | Volume | $\mu m^3$ | $V = \pi(l - w)(\frac{w}{2})^2 + \frac{4}{3}\pi(\frac{w}{2})^3$ |
| $F$ | Fluorescence | $a.u.$ | Total fluorescence intensity of the segmented pixels corresponding to A |
| $\tau$ | Division Time | min | Time elapsed since birth of a cell |
| $\frac{dF}{dt}$ | Production rate | a.u. $min^{-1}$ | Discrete derivative of fluorescence over time |
| $\frac{dV}{dt}$ | Growth speed | $\mu m^3 min^{-1}$ | Discrete derivative of volume over time |
| $\frac{1}{V}\frac{dF}{dt}$ | Specific production rate | a.u. $min^{-1}\mu m^{-3}$ | Production rate normalized by volume |
| $\frac{1}{V}\frac{dV}{dt}$ | Specific growth rate | $min^{-1}$ | Growth rate normalized by volume |

Table S1: Summary of key single cell measurements.

### Supplementary Figures

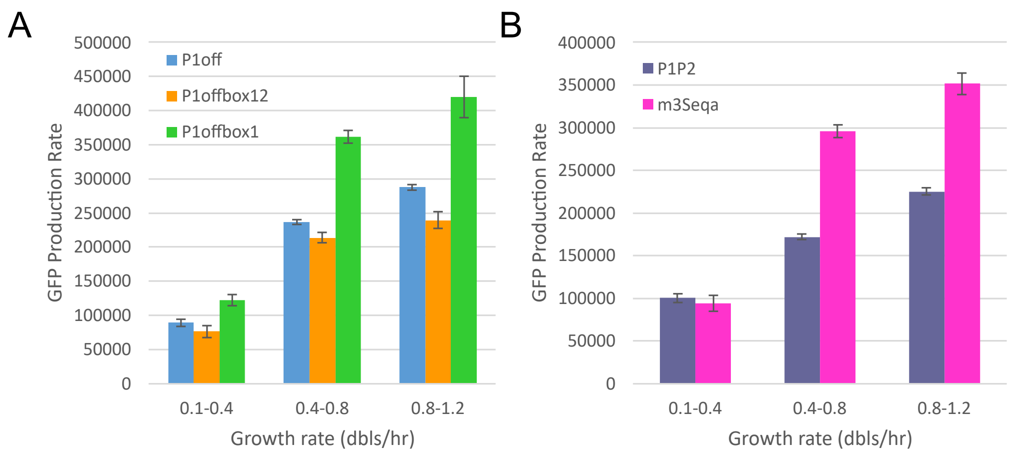


Figure S1: **Regulation by DnaA and SeqA at the population level.** The plots report plate-reader measurements of the change in population-average promoter activity for the different promoters in exponential growth phase at different ranges of growth rate, using the reporters described in Fig 1A of the main text. (**A**) Extent of activation and repression by DnaA of its own promoter. The effect of mutations of Box1 and Box2 agree with the previous data obtained on plasmids [19]. The decrease in binding affinity due to the mutation of Box1 results in an increase in the amount of DnaA-ATP required to bind at the promoter. The amount of DnaA-ATP in the cell is still sufficient for DnaA-ATP to bind to the higher affinity sites needed for activation but not to the lower affinity sites needed for repression. The result is an increased expression compared to the wild type. Mutation of Box2 results in a stronger disruption of DnaA binding and thus in a loss of transcription activation, seen as a decrease in expression compared to the Box1 mutant. (**B**) Extent of repression by SeqA of the dnaA promoter. The mutation of the GATC sites in the m3SeqA reporter system results in increased expression of the reporter protein, consistent with a decreased repression by SeqA. There is no effect of the same mutations in cells growing at a slower rate in M9-glu. It has been shown that a mutation in the SeqA gene or its binding sites at the dnaA promoter region has an effect mainly at faster growth rates [27]. The values shown are the average of three independent experiments. The error bar is the SEM.

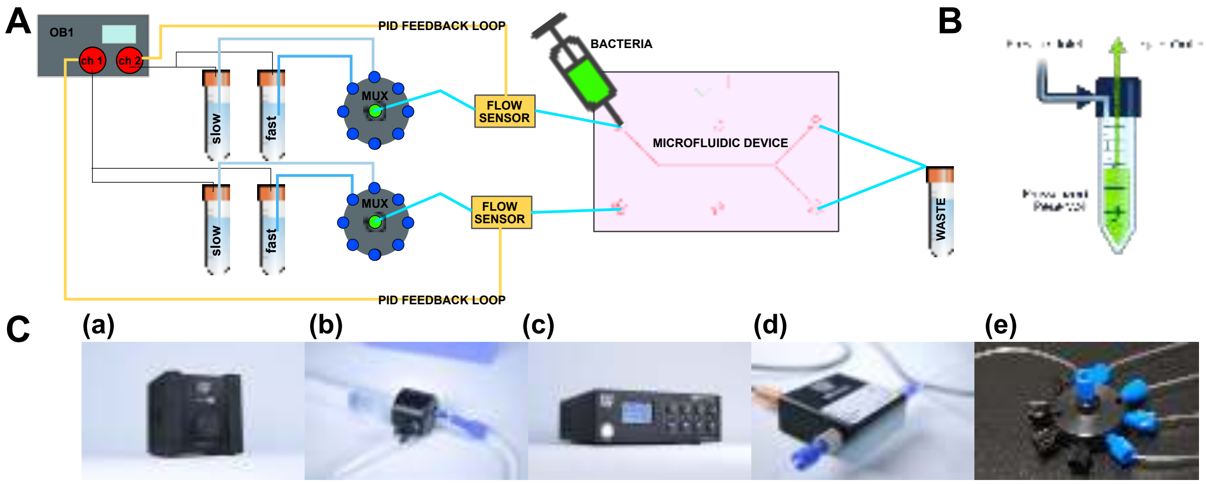


Figure S2: **Schematic representation of the microfluidic setup.** **(A)** The experimental setup used for upshift experiments (Elveflow). **(B)** Pressurized reservoir: the fluid flows through the outlet if gas pushes on the fluid surface. **(C)**, List of components:**(a)** Mux Distribution ,**(b)** Tubing, fittings and reservoirs ,**(c)** Flow controller OB1 Mk3+ , **(d)** flow sensor ,**(e)** manifold splitter 9 ports

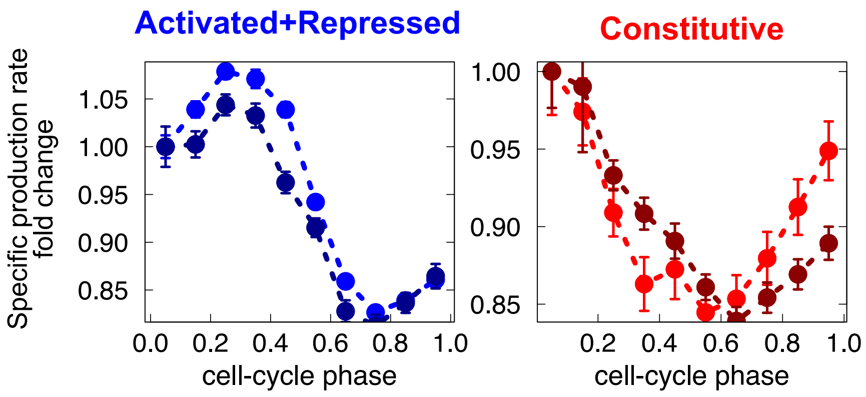


Figure S3: **Reproducibility across experimental replicates.** The plots report binned average of volume-specific production rates *versus* cell-cycle phase for the wt *dnaA*P2 (left) and constitutive promoter P5 (right) reporters from four distinct experiments, two replicates with each of the two strains. Each plot reports data from the two replicates using the same strain as circles with differently shaded colors. The dashed lines are guides to the eye. For both strains, different replicates show similar cell cycle-dependence of the conditional average of specific production rate at fixed cell-cycle phase.

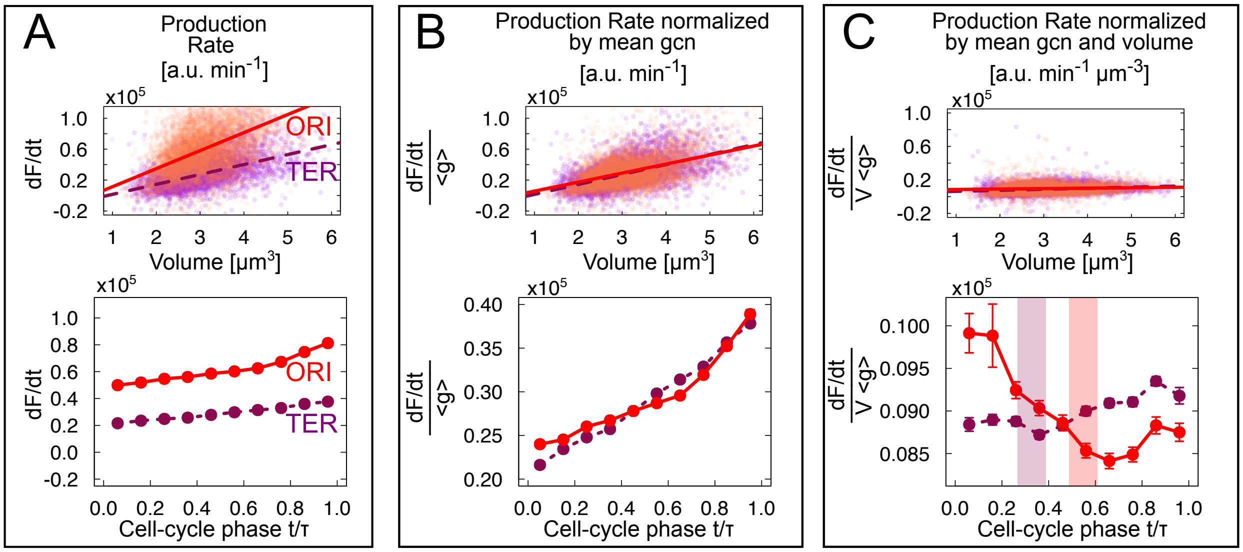


Figure S4: **The production rate of an unregulated promoter is proportional to cell volume and gene dosage.** **(A)** Production rate vs cell size (top panel) and vs cell cycle phase (bottom panel) for the P5 promoter inserted at origin-proximal (red, solid line) and terminus-proximal (purple, dashed line) loci. The production rate is proportional to cell volume and it is higher for the origin-proximal promoter, as expected from the higher gene copy number at the origin. This results in trends across the cell cycle. **(B)** Normalization by average gene copy number corrects the mean trend due to gene dosage in the same plots. **(C)** Normalization by volume and average gene copy number removes the main trends, but leaves the oscillations due to gene copy-number variation. This normalization shows small but significant cell cycle dependent differences between the production rates of origin-proximal and terminus-proximal unregulated promoters. The shaded areas correspond to the expected times of replication of the origin-proximal and terminus-proximal loci. Top panels (A-B-C) show scatter plot and trend lines (linear fits) of the quantities as a function of cell volume. Bottom panels (A-B-C) show binned averages of the quantities computed with respect to cell cycle phase. Error bars are standard errors of the mean from a re-sampled distribution of the signal, obtained by bootstrapping from the experimental data for each bin.

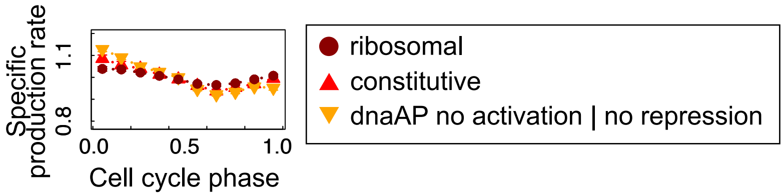


Figure S5: **A ribosomal promoter does not show oscillations beyond dosage.** As a control, we measured the expression of GFP from a minimal ribosomal promoter that also contains a GC-rich discriminator region (see ref. [42]). Average variations in volume-specific production of GFP from this promoter (circles) along the cell cycle are very similar to the *dnaA*P2 promoter stripped of all regulations (yellow inverted triangles) and to the non-ribosomal constitutive promoter (red upright triangles), indicating that the three reporters do not exhibit cell-cycle dependent oscillations beyond the effect of dosage.

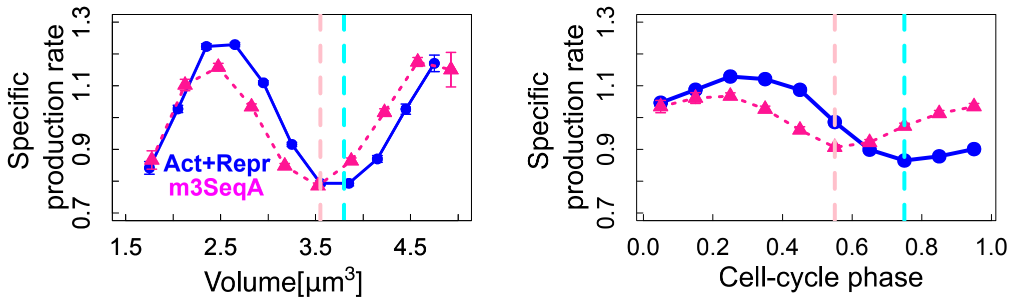


Figure S6: **The m3SeqA promoter, reporting for DnaA activity, shows shifted oscillations with a shorter period and smaller amplitude.** The plots are conditional averages of the volume and the specific GFP production change (plotted as fold change) as a function of cell cycle phase (left) or volume (right). Vertical lines show oscillation minima. The promoter repressed by DnaA-ATP presents oscillations within the cell cycle also when it is not regulated by SeqA (pink lines) although the timing of the peaks is altered and the amplitude of the oscillation is smaller. A mean volume at the minima around 3.5-3.6 $\mu m^3$ roughly corresponds to the expected mean cell volume at the mean cell-cycle phase (shown in Fig. S4C) corresponding to replication initiation in this condition.

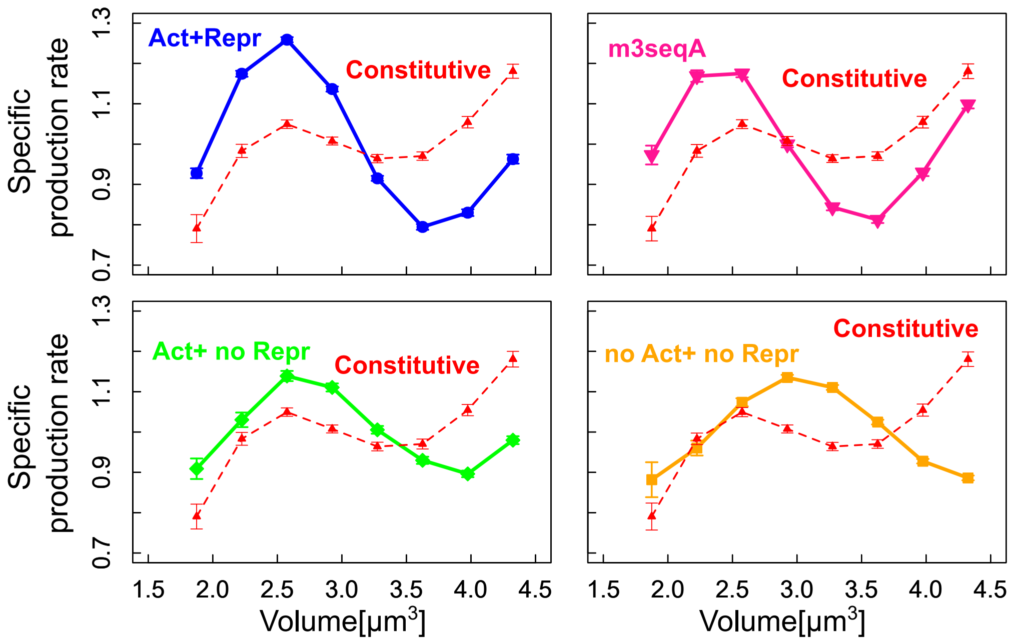


Figure S7: **Volume-specific GFP production rates show some oscillations with volume for all promoters**. Volume-specific GFP production rates show some oscillations with volume for all promoters including the constitutive promoter. The differences in amplitude and phase are consistent with SeqA delaying the increase in gene expression following gene duplication (comparison of no Act+noRepr with Constitutive), activation taking place after SeqA repression (compare Act+noRepr to noAct+noRepr) and DnaA-ATP repression taking place at a smaller volume than SeqA repression (compare Act+Repr to Act+noRepr). The plots show the fold change of conditional averages of specific production rate with respect to cell volume for different promoters.

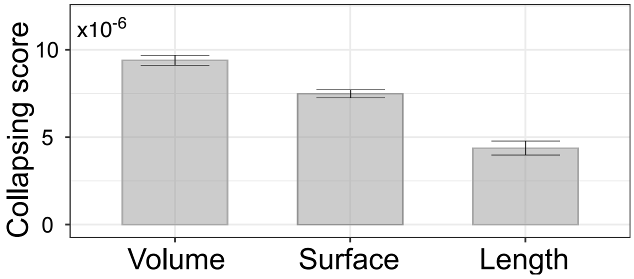


Figure S8: **Cell volume, not surface or length is the sensed variable.** The collapsing score defined in Fig 3 attempts to quantify direct sensing of a control variable from the independency from birth size of the oscillations of different conditional averages. In this plot the collapse score is computed considering volume, surface area and length possible control variables. The best collapse score is found by considering cell volume, suggesting that this is the control variable.

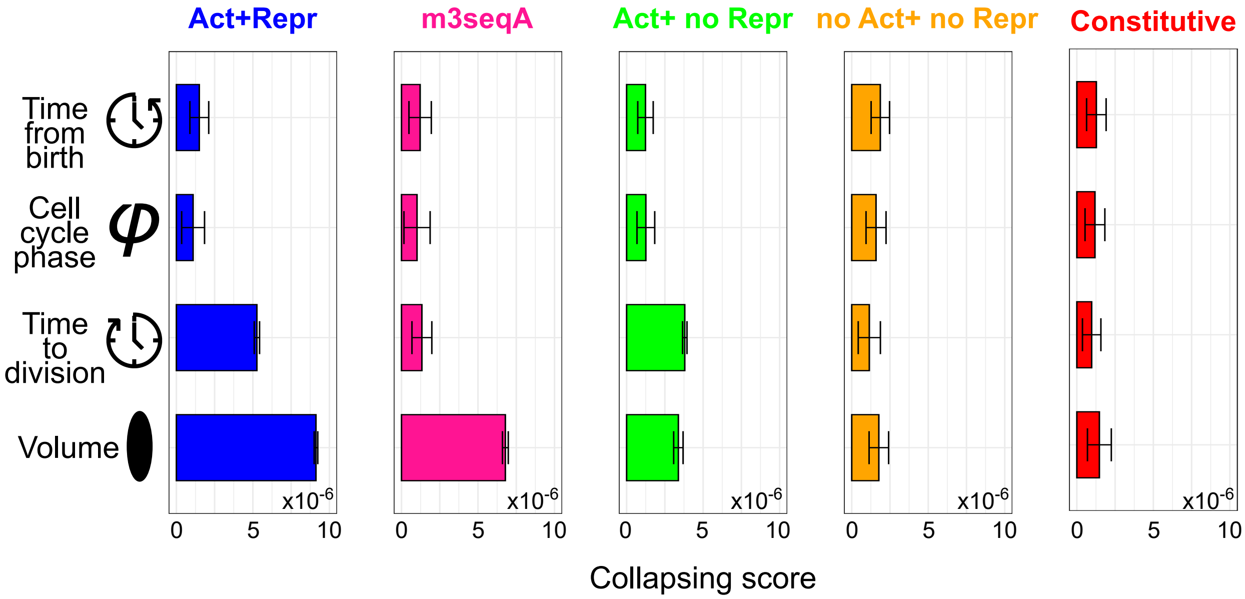


Figure S9: **Tests of direct volume sensing across different mutant promoters.** The plots show the collapse score of conditional averages of specific production rate for classes with different initial size (see Fig. 3 in the main text. The collapse score (see Fig 3) is defined as the SE-normalized inverse mean $L_2$ distance of conditional averages across bins of birth size.

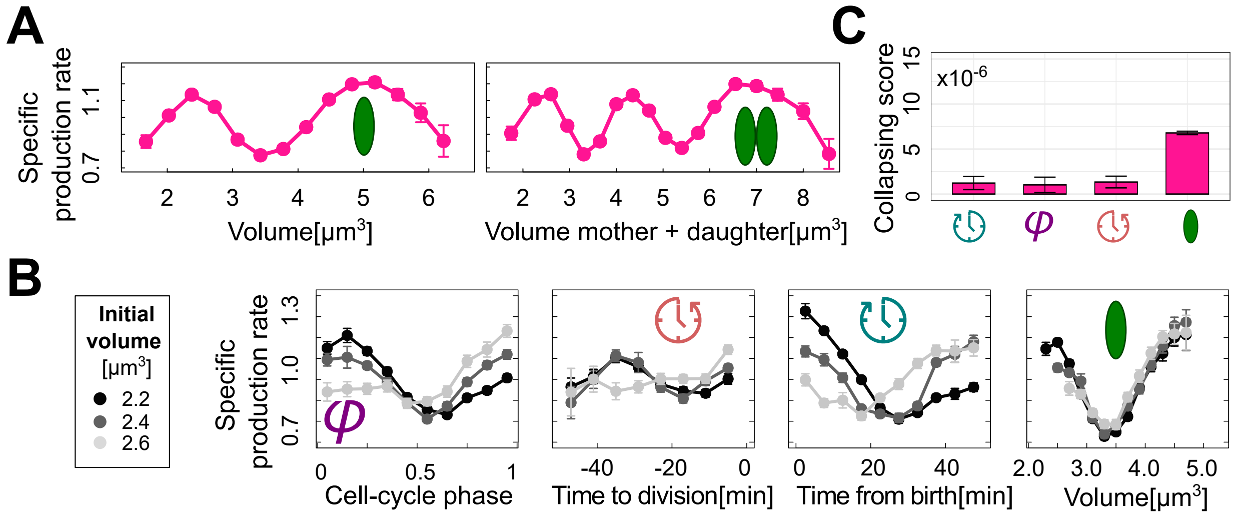


Figure S10: **Oscillations of the specific GFP production rate from the m3SeqA promoter, reporting for DnaA activity, collapse with volume when binned by cell size at birth but not with the time to division.** **(A)** (Left) Plot of the conditional average of GFP volume-specific production rate from the m3SeqA promoter as a function of cell volume. (Right) The same average across mother+daughter lineages shows two minima at multiples of a characteristic volume. **(B)** Volume-specific m3SeqA activity oscillations for cells of different initial size show different degrees of overlap when conditionally averaged as a function of cell cycle phase, time to division, time from birth and cell volume. The differently shaded curves result from data binned according to cell size (volume) at birth ($2.2 \pm 1\mu m^2$, black , $2.4 \pm 1\mu m^2$, dark-grey and $2.6 \pm 1\mu m^2$, light-grey). **(C)** Quantification of the collapse of the curves of panel B relative to 11 bins of cell sizes at birth (see Methods). Binning the data by cell size shows the best collapse. Error bars are standard errors of the mean obtained by bootstrapping from the experimental data for each bin.

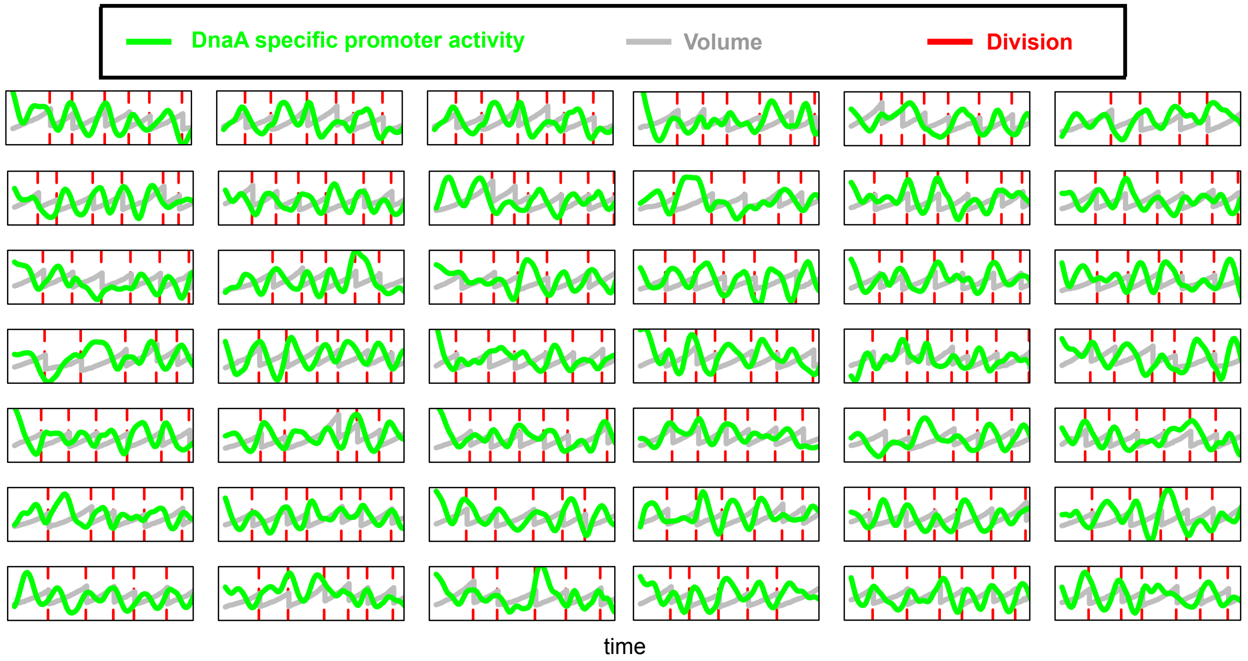


Figure S11: **Oscillations are observable at the single-cell level.** Set of lineages where volume (grey), *dna*AP2 promoter activity (green) and division (red) are tracked in single cells for several generations. The y axis is rescaled on order to show the two plots in the same range. The green lines refer to smoothed volume-specific derivatives of the fluorescence (see Methods). *dna*AP2 promoter activity shows strong oscillations that appear coordinated with cell-cycle progression.

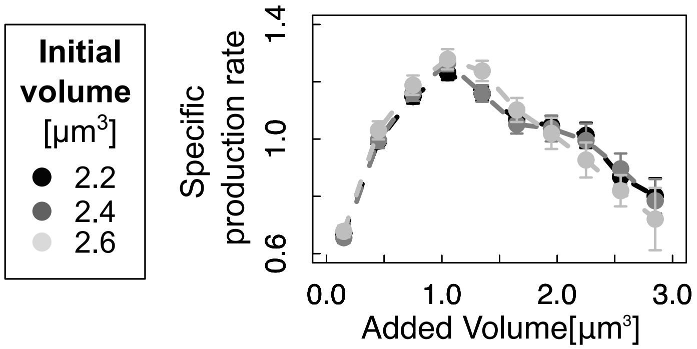


Figure S12: ***dnaA*P2 oscillations sense added volume.** Volume-specific *dna*AP2 promoter activity oscillations for cells of different initial size show a strong overlap when conditionally averaged as a function of added volume from oscillator minima. The differently shaded curves result from data binned according to cell size (volume) at birth ($2.2 \pm 1\mu m^2$, black, $2.4 \pm 1\mu m^2$, dark-grey and $2.6 \pm 1\mu m^2$, light-grey). If the variable in the $x$ axis is the sensed variable, the averages should change independently of the cell size at birth. Data for added volume show an equally good collapse as cell volume, hence this analysis cannot determine whether *dna*AP2 oscillator is a volume or added-volume sensor.

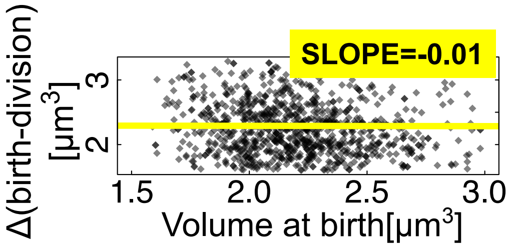


Figure S13: **Our experimental data confirm "adder" correlations between cell divisions.** The plot is a scatter plot of the added volume between birth and division versus volume at birth. The trend slope is compatible with zero, in line with an adder between divisions [55].

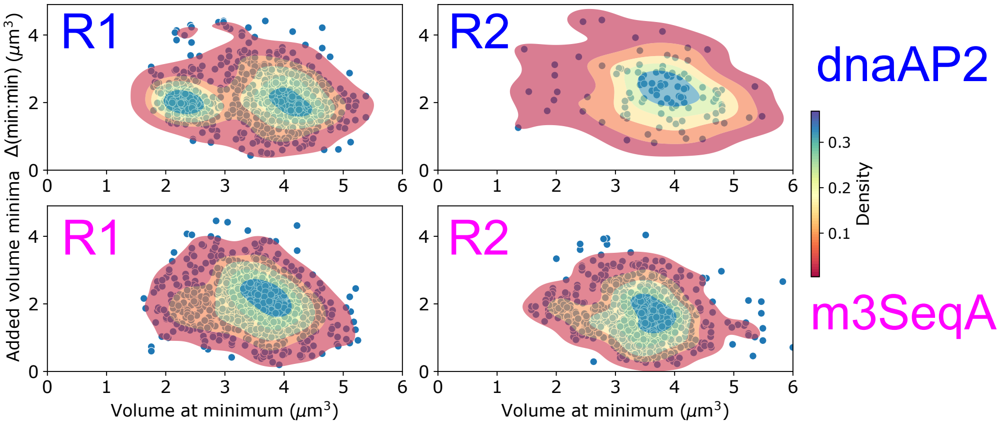


Figure S14: **Adder plots between *dnaA*P2 and m3SeqA oscillation minima differ.** The plots report the added volume between two consecutive minima separated by one cell division along wt *dnaA*P2 (top) and m3SeqA (bottom) lineages of more than 2 generations, *vs* volume at the first minimum, in two experimental replicates for each experiment. Scatterplots are shown superposed to density isolines obtained using a Gaussian kernel density estimator. The slopes of the main modes (isolated by a cutoff on the x-axis at 3 $\mu$m$^3$ for wt *dnaA*P2 and 2.1 $\mu$m$^3$ for m3seqA) evaluated by fits of the conditional averages are $-0.266 \pm 0.004$ (*dnaA*P2 R1), $-0.19 \pm 0.09$ (*dnaA*P2 R2), $-0.5 \pm 0.007$ (m3seqA R1) and $-0.18 \pm 0.04$ (m3seqA R2). The m3seqA R2 replicate yielded lower-quality time series and is likely more affected by noise in minima detection.

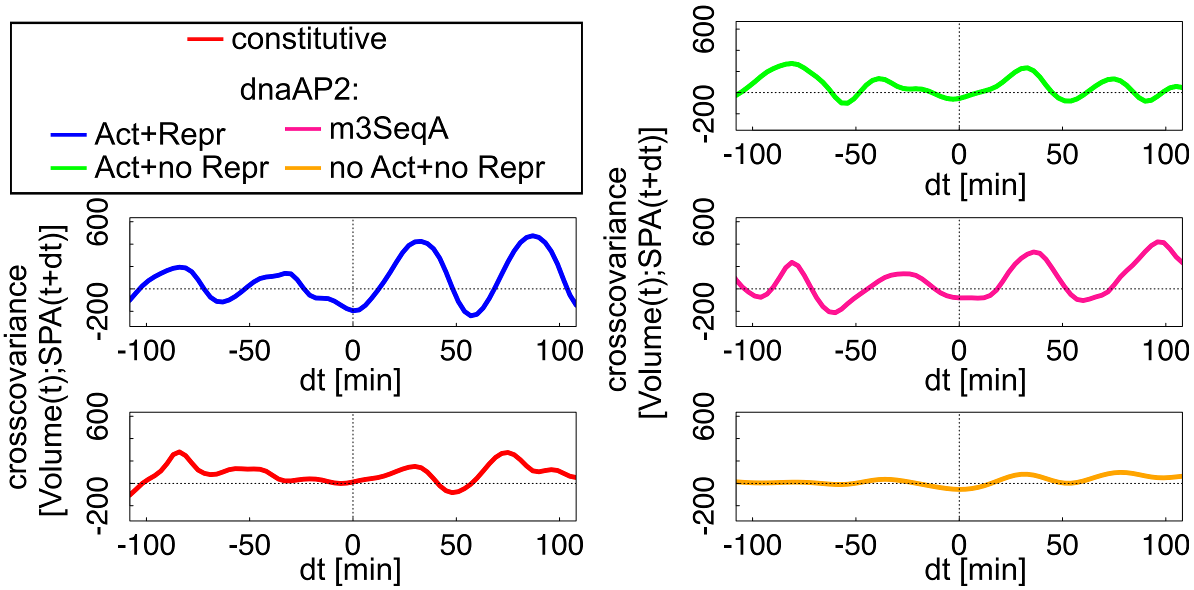


Figure S15: **Delayed cross-covariences show a volume-to-*dnaA*P2 time hierarchy for the *dnaA*P2 promoter.** The plots show delayed cross-covariance functions between volume time series and specific promoter activity for the different mutants of the *dnaA*P2 promoter. The function gives signatures of oscillations for all regulated promoters, but they are weaker for the promoters not regulated by DnaA or SeqA. Specifically, Of note, the m3seqA mutant gives a weaker signal than dnaAP2, supporting the idea that SeqA repression contributes to the correlation between the oscillator signal and cell division. The function is stringly asymmetric with respect to past *vs* future delays only for the wt *dnaA*P2 promoter, suggesting a time hierarchy between initiation-coupled oscillations and cell division.

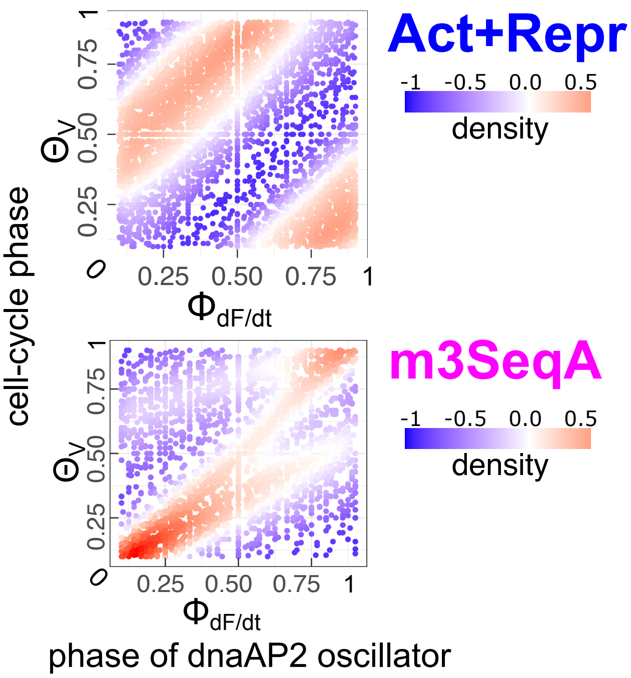


Figure S16: **The dnaAP2 oscillator is phase locked with cell-cycle progression, and the locking is lost by the m3SeqA promoter mutant.** The plots are a heatmap histogram of all the available lineage, using the trajectories of the phase of the *dnaA*P2 oscillator (x-axis defined as a variable that is zero at the oscillation minima and increases linearly to 1 at the next oscillation minima and the cell-cycle phase (y axis). The heatmap represents the so-called ”phase space” of the coupled oscillators. The wild-type data show that the heatmap is peaked at a well-defined value of the phase difference (top), a signature of a phase-locked synchronization state. This property is disrupted in the m3SeqA mutant, lacking the SeqA binding site (bottom). As the coupling depends on phase differences, the wild-type promoter follows the Adler equation for coupled oscillators.

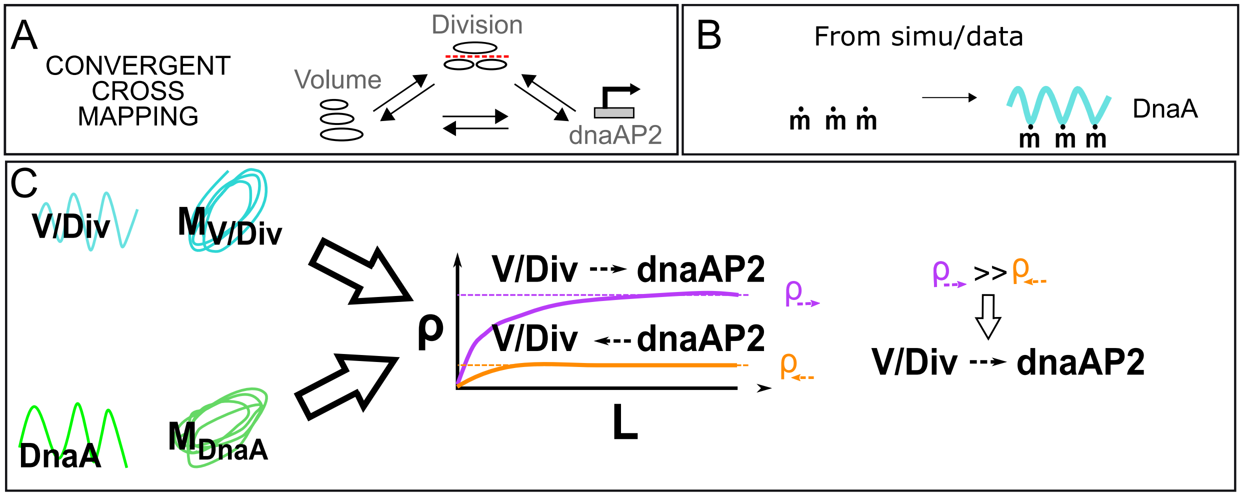


Figure S17: **Illustration of convergent cross-mapping (CCM) and its use with our data.** (**A**) We used CCM to detect causality between volume growth, division events and *dna*AP2 oscillations. For our experiments, these are time series directly obtained from lineage data along several generations. (**B**) In the case of experimental initiation data and simulated data we assume a cell-cycle oscillator made by sinusoidal oscillations with amplitude 1 whose minima are placed at initiations. (**C**) Explanation of CCM. CCM looks for the signature of causal effects of variable $A$ in variable $B$'s time series by seeing whether there is a correspondence between the "library" of points in the attractor manifold built from $B$, $M_B$ , and the points in the reconstructed attractor manifold for variable $A$, $M_A$. The two manifolds are reconstructed from lagged coordinates of the time-series, and $L$ sample size used to construct the library (which also describes the size of the delay time windows that are used for prediction) setting the dimensionality of the manifold embedding. The accuracy of predictions varies as a function of embedding dimension $L$, and should plateau for sufficiently large $L$. The *a priori* asymmetric coefficients $\rho_{AB}$ and $\rho_{BA}$ are conditional correlations between the two variables at fixed coordinates along the manifold of one of the two, and can be interpreted as causality coefficients. One can estimate an optimal embedding dimension by using simplex projection to test the ability of a process to predict its own dynamics through leave-one-out cross-validation.

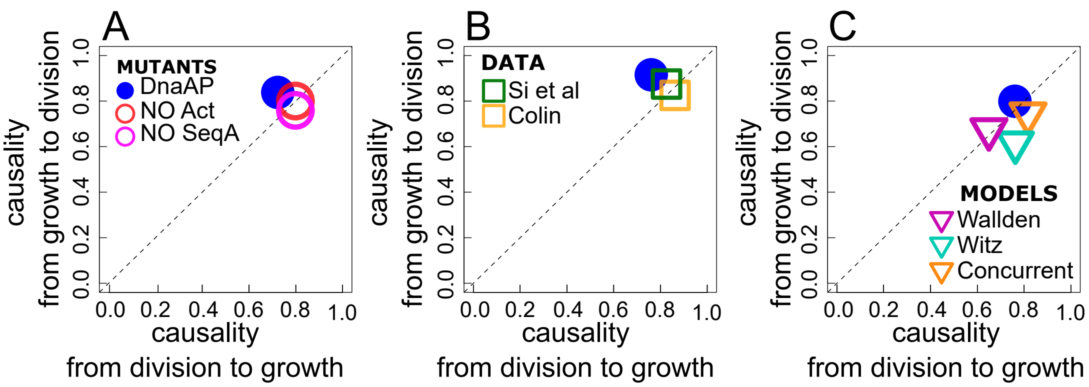


Figure S18: **Convergent Cross-Mapping (CCM) scores the cell-volume and the cell-division oscillators as causally equivalent.** (**A**) The CCM causality coefficients are symmetric between the volume time series and a time series constructed by pulses of 10 minutes around cell division events. The three plots refer to our experimental data, experimental data from the literature and simulations of common cell division models from the literature (See Fig. S17 in the main text). The symmetry suggests a mutual coordination between the two time series.

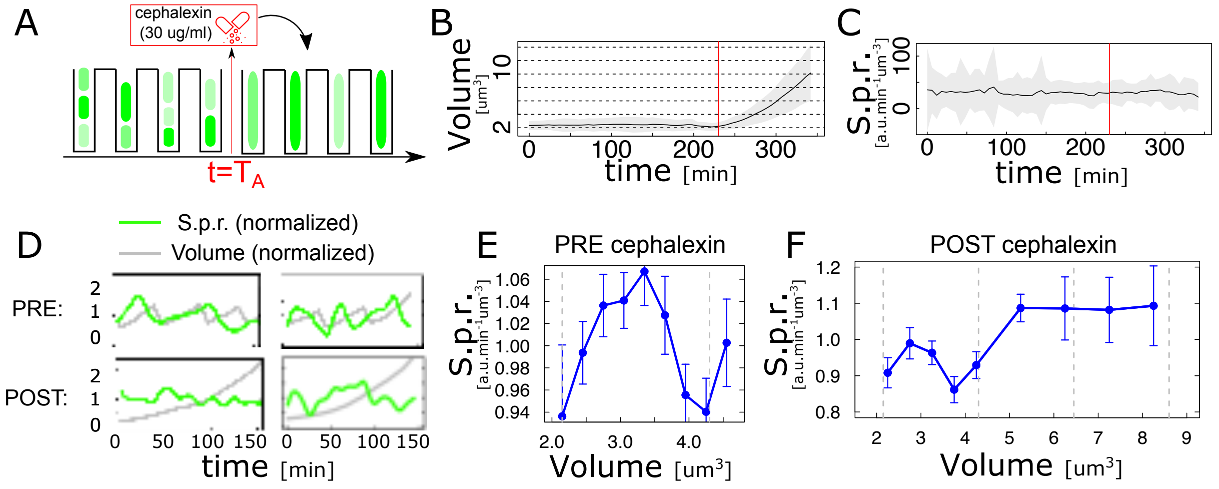


Figure S19: **Cephalexin inhibits division, affecting *dnaA*P2 oscillations**. When Cephalexin is added, cells cease to divide (**A**) and volume keeps increasing over time (**B**). Panel **C** *dnaA*P2 specific production rate remains constant when averaged conditionally to time elapsed from the perturbation. Hence, the oscillations are not synchronous with respect to this variable. (**D**) *dnaA*P2 keeps oscillating upon cephalexin treatment. The plots show *dnaA*P2 specific production rate and volume at single cell level with (top) and without (bottom) cephalexin. (**E**) Specific production rate conditionally averaged as a function of cell volume without antibiotic shows strong oscillations, which are lost (**F**) after the first cell cycle upon treatment with cephalexin. Hence cephalexin treatment appears to destroy the synchronisation of *dnaA*P2 oscillations with cell volume.

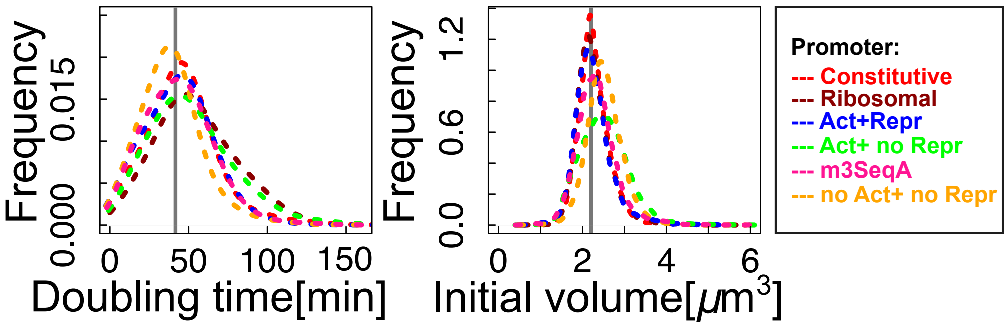


Figure S20: **Doubling-time and birth-size distributions are consistent for strains carrying different reporters.** The plots show normalised histograms for the two quantities taken over around 3000 cells for each strain carrying different GFP reporters. The distribution of interdivision times (left) is centered around 45 minutes (dark grey bar) for all strains, while the distribution of cell birth size (right) is centered around 2.1 $\mu m^3$ (dark grey bar).
